# MASLD-Bench evaluating language models on liver omics and fibrosis endpoints

**DOI:** 10.64898/2026.11.20.768002

**Authors:** Subhadip Banerjee, Wim Vanden Berghe, Rawiwan Charoensub

## Abstract

Language models may provide an interface to molecular measurements, but their ability to predict fibrosis remains uncertain. MASLD-Bench retrospectively evaluates archived model outputs from fixed liver-omics datasets. We examined 19 task datasets containing 106,688 prompts across liver transcriptomics, staged cohorts, progression, perturbation transfer and plasma cell-free RNA. Evaluated subsets comprised 15–414 items per model–task combination from 21–179 samples per family. Metrics included accuracy, mean absolute error (MAE), response-parsing coverage, 95% confidence intervals, paired differences and exact McNemar tests.

The tissue analysis included 17 eligible models. On 163 common binary items, six matched majority accuracy (0.656); none exceeded it. Nemotron’s five-class accuracy was 0.276 versus majority accuracy of 0.264 and uniform-choice chance of 0.200. Regression MAEs among parsed responses were 1.142 for Qwen3-1.7B v5 (162/162 parsed) and 1.517 for Llama 3.3 70B (150/162 parsed); corresponding MAEs on 150 jointly parsed items were 1.127 and 1.517. Rankings varied across formats and populations. In the second plasma cell-free RNA cohort, logistic regression achieved accuracy of 0.609 (95% CI, 0.536–0.676) with training-only per-gene preprocessing, versus majority accuracy of 0.531. With per-sample preprocessing, the interval included the majority rate. A five-gene panel selected in the reference cohort achieved external accuracy of 0.615 without established paired superiority over the majority classifier (McNemar p = 0.133).

Baseline definition, training exposure, output validity and preprocessing affect interpretation. These retrospective findings establish neither clinical utility nor a general language-model advantage in fibrosis staging.

**Plain Language Summary:** Artificial intelligence (AI) systems, including large language models, are increasingly being tested to see if they can read complex medical data and help predict disease. In this study, we evaluated several popular AI systems to see if they could predict liver scarring (fibrosis) in patients with MASLD, a common chronic liver disease. We presented the AI models with gene activity data from liver biopsies and blood samples and asked them to stage the disease.

We found that none of the AI models outperformed a simple "majority rule" — meaning no model was more accurate than simply predicting the most common outcome seen across all patients. While some models appeared to have very low error rates, this was misleading: the models were simply guessing the average patient response rather than actually learning meaningful biological patterns from the data. Our study also discovered that some models had inadvertently been trained on the very test data they were being evaluated on, which would artificially inflate their apparent performance. Additionally, the evaluation software sometimes cut off the AI’s responses before they could be completed, which could affect the results.

Our findings indicate that much stricter and fairer testing is needed before AI systems can be trusted to interpret molecular data or assist doctors in diagnosing liver disease. Rigorous evaluation that prevents training on test data and captures complete model responses is essential before AI can play a reliable role in clinical medicine.

## 1. Introduction

Metabolic dysfunction-associated steatotic liver disease (MASLD) affects more than 30% of adults worldwide and is a leading cause of chronic liver disease [1,2]. Histological staging — fibrosis stage F0–F4, steatohepatitis activity, ballooning — remains the reference standard against which non-invasive tests are judged [3,4], and those non-invasive alternatives remain imperfectly calibrated to histology [5]. Widely used blood-based scores such as the Fibrosis-4 index (FIB-4) and the NAFLD Fibrosis Score (NFS) leave a substantial indeterminate range, which is one reason the field continues to look for better molecular readouts.

Large language models are being investigated as tools for interpreting biomedical information. Molecular datasets, however, contain numerical measurements and ordered outcomes that depend on expert assessment. MASLD therefore provides a useful setting in which to ask whether a language model can predict a histological label from a fixed panel of gene-expression values presented as text.

Related evidence spans from biological benchmarks and clinical language-model studies. LongevityBench evaluated 17 tasks across five biodata domains and 18 language models, and reported zero-shot performance at or near chance on omics pairwise tasks (GTEx transcriptomic age 0.48–0.54; proteomics near-random), while the same systems succeeded on clinical and tabular tasks (NHANES 10-year survival 0.882; SynergyAge 0.781) [6]. The study reported that compact fine-tuned "Longevity-LLMs" matched or exceeded API-accessed models at far lower inference cost [6,7]. Clinical studies also investigate language models for liver-disease assessment and risk stratification [8,9]. In liver disease specifically, Fibro-GPT reported AUROC 0.91/0.90 for advanced fibrosis, but its inputs were seven routine laboratory values rendered as text, its labels were a FIB-4/NFS+VCTE surrogate rather than histology, its cohort numbered 162, and thresholds were selected on the same cohort that produced the reported metrics [10,11]. Pugliese et al. discuss opportunities for generative AI in MASLD [12]. A separate study used language models to extract structured information from radiology, pathology and transplant-assessment reports and reconstruct longitudinal clinical trajectories in chronic liver disease [13]. These clinical-text tasks differ from predicting fibrosis from numerical expression panels and motivate evaluation of liver *gene-expression* inputs against *biopsy*-based outcomes.

MASLD-Bench focuses on instruction-following models given fixed molecular panels and fibrosis endpoints. Related resources evaluate single-cell integration, perturbation prediction, bioinformatics agents and omics discovery workflows [14,15,16,17]. Our evaluation asks a different, bounded question and does not claim priority over all omics benchmarks. For binary classification, paired comparisons on a common test set can use McNemar’s test to evaluate differences in error rates [18]. We therefore align evaluated items before comparing models and distinguish descriptive rankings from statistical comparisons; statistically grounded language-model evaluation provides related context [19]. Reviews of artificial intelligence in liver-disease omics motivate this assessment [20].

The benchmark records source cohorts, answer formats, evaluated populations and preprocessing procedures. A frozen corpus is a versioned, checksum-identified snapshot: it fixes the evaluated bytes but does not establish prospective registration or eliminate analytic flexibility. We distinguish construction counts from independent samples and disclose training exposure before interpreting held-out performance.

Two further design choices bound what the benchmark can say. First, the tested inputs are molecular measurements only — expression panels rendered as text. Routine clinical covariates (age, sex, BMI, aminotransferases, platelets, HOMA-IR) and clinical risk scores (FIB-4, APRI, NFS) are deliberately excluded. This makes the comparison a test of molecular readout rather than a comparison against clinical practice, and it means the benchmark cannot establish superiority over FIB-4, APRI or NFS. Second, the benchmark evaluates models as they are, without new training for the API-accessed model group; the compact arm is fine-tuned on a declared mixture. Differences between arms therefore confound model class with training exposure, which is why transfer experiments require separately verified evaluation procedures rather than treating the fine-tuned arm as a clean scale comparison.

We report a retrospective analysis of the frozen archives. The contribution is an auditable evaluation of molecular-input language models that exposes the effects of baseline definition, sample overlap, output validity and preprocessing on interpretation. It is not a new learning algorithm or a prospective clinical validation.

## 2. Methods

### 2.1. Benchmark datasets

To create MASLD-Bench, we constructed structured prompts from publicly available cohorts and one plasma resource accessed under a Data-Usage Agreement. Each source was prepared in one or more answer formats — binary classification, five-class classification, ordinal regression, pairwise comparison or numeric regression — and saved as a separate, versioned dataset before model evaluation. Features were selected without using outcome labels and presented as identifier–value pairs with two decimal places. Normalisation and standardisation are described below; the cfRNA datasets use per-gene standardisation across samples. The complete manifest (v3-freeze-17) records the processing rule for each dataset, item counts, checksums for the datasets and source matrices, and the global seed (20260929). Corrections produce a new version, retain superseded archives separately and add a dated entry to the run log. The freeze-6 and freeze-16 corrections are reported here. The OSF record is a retrospective deposit of the analysis plans and run log; version dates alone do not establish prospective registration. This frozen dataset is a versioned, checksum-identified snapshot that permanently fixes the exact data matrices and text prompts presented to the models. This practice ensures that the evaluated inputs can be audited and reproduced without risk of silent dataset updates. However, we explicitly distinguish this data-level freezing from prospective registration. Whereas registration requires the pre-specification of hypotheses and statistical analysis plans before data access, freezing only guarantees input data integrity. It does not retroactively certify that the analysis scripts were written without knowledge of the outcomes, nor does it eliminate researcher degrees of freedom during statistical evaluation

### Liver transcriptomics (MB-20xx)

The core evaluated subset is the EPoS biopsy cohort — the 168 fibrosis-staged samples of GSE135251 described by Govaere et al. [21], obtained as the re-distributed count matrix of the Kamzolas et al. [22] supplementary deposit. The frozen test corpus contains 168 of the 216 GSE135251 samples and is rendered as 11,653 prompts covering binary advanced fibrosis (F3–F4 versus F0–F2), five-class stage (F0– F4), ordinal stage regression and intra-cohort pairwise comparison. The pairwise task presents two profiles and asks which has the more advanced stage. All 168 frozen sample identifiers matched GSE135251 sample records. A separate expression-level spot check compared prompt values for GSM3998167 with raw counts for that same sample and found Pearson r = 0.999996. This correlation was computed for one sample, not for all 168 matched samples; it supports accession attribution but does not establish a cohort-wide linear transformation or bit-for-bit identity. The identifier comparison and the expression-level spot check are distinct pieces of evidence. Neither recovers the complete sequence of source files and processing steps, and the expression check should not be interpreted as validation of every frozen sample profile.

### Train–test overlap disclosure (MB-20xx / MB-12xx)

The staged-liver SFT pool described below was built from the same GSE135251 series: all 168 MB-20xx test samples are also present in MB-12xx, and the v6/v7 checkpoint mixtures upsample MB-12xx four-fold. The checkpoints Qwen3-0.6B v6, Qwen3-1.7B v6 and Qwen3-8B v7 therefore saw every tissue test sample during supervised fine-tuning, and their tissue results are not held-out evaluation. In addition, the MB-12xx binary task uses an F2–F4 threshold for "advanced fibrosis" while the frozen MB-2001 test uses F3–F4, so the SFT data taught the opposite binary label for 42 of the 168 shared samples (the F2 samples); those 42 disagreements are a cross-task label-definition difference and are not evidence of a labelling error. The five-class and regression labels agree between the two datasets. Earlier checkpoints (v3, v5) were not exposed through the documented project fine-tuning mixtures. API-accessed models’ pretraining content is not established here. The primary tissue analysis therefore excludes the three exposed checkpoints and reports them separately.

### Scope of the overlap check

An all-corpus comparison of 11 training files against 15 test files found the exact-ID overlaps listed above (MB-20xx and the five MB-40xx geometric/k-shot test datasets) and no additional exact-ID overlaps within the inspected comparisons. This is not a universal patient-identity exclusion: some datasets lack donor aliases, and normalized numeric identifiers from unrelated naming schemes were not treated as proof of identity. No v6/v7 MB-40xx response archive was found, so the exposed checkpoints have no reported MB-40xx result to interpret.

### Staged-liver training pool (MB-12xx)

Three public biopsy cohorts (GSE135251, n = 216; GSE167523, n = 98; GSE130970, n = 78) were harmonised to a common set of genes and rendered as 1,196 training prompts across stage, activity and ordinal formats. A fourth candidate cohort was obtained but excluded: its series matrix carried no per-sample numeric key that could be joined to the expression columns, and order-matching was rejected after the freeze-6 correction.

### Progression proxy (MB-85xx)

GSE185051 contributed 104 prompts from 52 paediatric NAFLD patients and five controls, staged F0–F4. Because no paired-biopsy longitudinal data were available, progression is inferred cross-sectionally from fibrosis stage; only five of 52 patients are progressive. The family therefore functions as a negative control and was plan-specified as one.

### Blood-proxy severity (MB-86xx)

GSE267032 contributed 57 prompts from 21 patients with a three-level whole-blood severity grade (8 low, 6 medium, 7 high) in three formats. The corpus was rebuilt at freeze-16 after a one-row header misalignment. Controls in the deposit contained no expression measurements and were excluded.

### Liver-to-plasma proteomics (MB-87xx)

MSV000094959 contributed 422 prompts in two formats: a three-class platform-identity task, which measures batch separation and is reported as such, never as clinical signal; and a masked-protein regression task in which the 30 proteins with the highest variance are selected per study, one is masked in rotation, and the model predicts its z-score from the remaining 29.

### Primary-hepatocyte perturbation (MB-88xx)

GSE247407 contributed 30 prompts from five donor preparations, each with three replicates under control and free-fatty-acid exposure; the sample key is the donor, so grouped cross-validation blocks donor leakage by construction. A separate training pool from E-MTAB-15601 (HepG2 DRUG-seq, 40 compounds plus DMSO, 992 prompts) supplies the cross-system transfer experiment: train on HepG2 compounds, test on primary hepatocytes.

### Plasma cell-free RNA (MB-89xx)

The Ghent plasma cfRNA resource (S-BSST1614; matrices for cohorts 1 and 2, annotations from Zenodo 10.5281/zenodo.19047046) was joined to labels with explicit key verification (112/112 and 821/821). Three datasets were frozen.

The reference cohort contains 336 prompts from 112 samples with Brunt fibrosis stages F0–F4 assessed by a UZA biopsy reader and an indicator of agreement between readers for each sample. An independent AMC reader assessed 108 of these samples. The AMC dataset contains 324 prompts from those 108 samples; the prompt text is identical to that used for the corresponding reference-cohort samples, so only the reference labels differ. The second cfRNA cohort contains 537 prompts from 179 patients, with one sample selected per patient, clinical-consensus reference labels and biopsy assessments for a 42-patient subset. The first Baseline row is selected for each patient, falling back to the first available row. This rule depends on row order and is a limitation.

### Rendering

For all families, expression panels are rendered as identifier–value pairs (Ensembl identifiers, or gene symbols where the source is symbol-keyed). Normalisation is label-free: counts per million, then log1p. Panels are the top 200 genes by median absolute deviation, computed internally to each corpus.

### Preprocessing procedures

The frozen MB-89xx datasets standardise each gene across samples. The top-200 median-absolute-deviation panel and per-gene means and standard deviations are estimated on the full corpus, so classical cross-validation on this representation is transductive: held-out samples influence the representation. Two sensitivities select the panel within each training fold: one estimates per-gene means and standard deviations on training samples only, and one standardises each sample across its selected genes. These analyses do not regenerate frozen LLM prompts. Classical baselines reconstruct unrounded matrix values, whereas prompt text carries two decimal places; they share panels and samples but not identical numerical inputs.

### 2.2. Language model benchmarking

We evaluated API-accessed models through a single router, archiving the requested model string per response: Nova Pro, Claude Sonnet 4.5, DeepSeek Chat, Gemini 2.5 Pro, Llama 3.3 70B, Nemotron 120B, GPT-4.1, GPT-4.1-mini, GPT-4o-mini, GPT-5.6 Luna, Qwen2.5 72B and Qwen3 235B. Decoding was zero-shot, single pass, temperature 0.0, top-p 1.0, with a 1,500-token cap in the analysis plan and a 900-token cap in the cfRNA evaluation script actually used. We report the executed budget, not the planned one. Compact-model evaluation used a 24-token output budget, which is relevant to the observed truncation and parsing failures.

### Model inclusion

The twelve named API systems constitute the cfRNA model set. The tissue table additionally includes the archived Inception Mercury-2.5 system (identifier inception mercury-2.5). Thus the 17-model eligible tissue model set comprises 13 API systems and four v3/v5 compact checkpoints. The complete tissue model set has three additional exposed v6/v7 checkpoints.

Two evaluation-software corrections are material. The first API-model evaluation pass capped output at 200 tokens, truncating verbose reasoners before their verdict; a 900-token re-pass of unparsed cells plus two parser fixes recovered approximately 750 records at no additional API cost. Both passes are archived. A parser revision is a scoring change, not a model change, and is reported as such.

### Operational failure versus model error

The evaluation script archives the requested model name, the raw text, whether a parser produced a value, and any API request failure. It does not archive a provider-returned model identifier, usage counters or a finish reason. A requested name therefore does not prove a frozen provider snapshot, and this is a provenance limitation. Where the archive contains API request failures, those rows are classified separately from model answers: an HTTP failure is an operational failure of the evaluation, not evidence that the model lacks a capability. This distinction is not cosmetic. One of the API-accessed models returned HTTP 404: model unavailable on all 873 cfRNA requests and produced zero valid completions; its zero accuracy is reported as a failed evaluation, not as a measurement.

### 2.3. Classical baselines and paired statistics

Classical binary analyses use logistic regression and histogram gradient boosting. The ordinal cfRNA analyses use Ridge regression and a histogram-gradient-boosting regressor. Cross-validation uses five folds, with stratification and grouping as specified for each task. Reported bootstrap intervals use 2,000 resamples and fixed seeds. Unless explicitly stated otherwise, intervals resample fixed predictions rather than refitting the complete model and selection procedure in each bootstrap replicate; they do not capture all training or selection uncertainty.

### Binary comparison criterion

This criterion compares the lower bound of a 95% bootstrap accuracy interval with the observed binary majority rate in the evaluated labels. This test-set majority is a descriptive reference, not a training-estimated classifier. Meeting this confidence-interval criterion is distinct from establishing paired superiority against a constant classifier on the same items.

### Ordinal comparison criterion

Each regressor is compared with a constant predictor fitted as the median label in each training fold. We report the paired difference in absolute error, defined as constant-predictor error minus model error. An interval spanning zero does not demonstrate improvement; a positive point estimate alone is insufficient.

### Multiplicity

Where more than one comparison supports the same conclusion, the family is declared and Holm-adjusted p-values are reported. Selection of the best model on the evaluation set is disclosed wherever it happens. Prompt-level bootstrap intervals treat prompts as independent, which is liable to underestimate uncertainty for repeated donors and for pairwise formats that re-use the same profiles; we state this rather than presenting the intervals as cluster-robust.

### Equivalence

A non-significant superiority test is reported as "no statistically detectable difference". No equivalence or non-inferiority margin was pre-specified, so no equivalence claim is made anywhere in this paper.

### Eligible-model tissue analysis and evaluated populations

The primary tissue model set is every model with all five tissue tasks, excluding the three checkpoints exposed to MB-12xx. Single-sample tasks are scored on the intersection of archived (task, sample) keys across that model set; absence from an archive is recorded as "not archived" and is not silently scored as an error. Pairwise results are grouped by the exact set of queried item keys and are never averaged across different evaluation sets. The response archives do not store prompt text, so prompt-byte identity cannot be verified from them; the validated join is task, sample identifier and reference label. Classification uses invalid-as-error. Regression reports MAE among parsed numerical responses with valid/n, an in-domain sensitivity analysis (0–4 for fibrosis, 0–8 for the NAFLD Activity Score (NAS)) and an MAE with penalties for missing or out-of-range predictions. The latter uses absolute error for in-domain predictions and assigns a penalty of 4 (fibrosis) or 8 (NAS) to unparsed outputs or predictions outside the permitted score range, averaged over all evaluated items; it is a declared sensitivity metric, not an estimate of the missing answers. Pairwise intervals use a 2,000-replicate biological-sample cluster bootstrap; single-sample intervals use item-level bootstrap. Ranks are computed within the same model set and item set, and the cross-format mean rank excludes pairwise because the evaluated item sets differ.

### 2.4. Compact model training

The smaller-model series contains Qwen3-0.6B and Qwen3-1.7B full-LoRA runs under three training configurations (v3, v5, v6) and one Qwen3-8B quantised LoRA run (v7, 4-bit NF4). v3 introduced format-aware answer construction and class-balanced upsampling; v5 expanded the mixture; v6 added the 40-compound DRUG-seq pool and the staged-liver pool (48,221 training examples); v7 applied the v6 training configuration to the freeze-15 datasets (49,157 examples).

### Definitions used throughout

Compact models are the evaluated 0.6B–8B open-weight Qwen3 models. API-accessed models are the systems evaluated through the named API services. This group includes both proprietary and open-weight models and is not restricted to models with at least 70B parameters. These labels describe the comparison groups, not a uniform difference in model size, licensing or capability.

### 2.5. Metrics, definitions and reporting policy

We define each metric at first use and report only quantities that were actually computed for the stated model, task, label source and sample set.

- **Accuracy** — proportion of items answered correctly. Invalid or missing answers count as errors in the headline figure.

- **Sensitivity** — true positives divided by all items whose reference label is positive, including positives the model failed to answer. The denominator is always stated.

- **Specificity** — true negatives divided by all reference-negative items, same convention.

- **Balanced accuracy** — the mean of sensitivity and specificity. A complete, valid always-negative predictor scores 0.5 only when every item receives an answer. Under the invalid-as-error policy used here, unparsed items count against the relevant class denominator, so an always-negative model that also fails to answer some items scores below 0.5: the reference cohort GPT-4o-mini result is 0.46875, not 0.5.

- **AUROC** — probability that a randomly chosen positive item is scored above a randomly chosen negative item; tie-aware, computed with average ranks.

- **Average precision (AP)** — the step-interpolated area under the precision–recall curve, computed with a tie-aware implementation. AP and trapezoidal PR area are different statistical quantities and are not used interchangeably.

- **Mean absolute error (MAE)** — mean absolute difference between predicted and reference values. Reported conditionally, over parsed answers only, with the valid n and coverage stated; an all-item failure-aware figure is reported alongside where a penalty policy is declared.

- **Spearman ρ** — rank correlation between predictions and reference labels.

- **Cohen’s κ** — agreement beyond chance. Unweighted κ is reported for nominal categories and quadratic-weighted κ for ordinal stages, with denominators stated.

### Rounding

Proportions and interval bounds are generally displayed to three decimals; criterion decisions use the unrounded values in the analysis artifacts.

### Cohort terminology and prediction variability

The reference cohort is the 112-sample cfRNA cohort; it supplies the development data for the five-gene panel. The second cfRNA cohort contains 179 patients and is used for external evaluation of that panel. Separately, reduced prediction variability means that predicted stages vary less than the observed stages. Low variability and weak rank correlation do not establish that a model predicts the training-set mean. We therefore report prediction means and standard deviations alongside those of the observed stages, without assigning a mechanism to this pattern.

### Distinguishing analysis types

Three cfRNA analyses are reported separately and are not inferential tests: post-hoc marker overlap against a deposited cell-type Gini table; a top-k panel analysis with feature selection nested inside cross-validation; and discrimination rescoring of frozen ordinal outputs. None of them generates a p-value that supports a causal claim, and a classical-model ablation cannot establish which features a language model used.

### Canary probes

A canary is a deliberately planted unique sequence used to test whether a model has memorised training data. No canary was planted in these datasets. A probe planted now could not retrospectively exclude contamination in archives collected earlier, so memorisation testing is stated as future work.

### 2.6 Software and Dependencies

Computational analyses were implemented in Python (v3.10+). Data operations utilized pandas, numpy, csv, json, pathlib, and ast. Statistical analyses (bootstrap CIs, McNemar’s test, Mann-Whitney U, Pearson r) were conducted using scipy and numpy. Figures were generated with matplotlib and seaborn and saved as high-resolution (300 dpi) PNG and PDF files.

### Reproducibility

To ensure environmental and data integrity, exact package versions are pinned in requirements-release.txt, and SHA-256 hashes for every file in the submission package are recorded in RELEASE_MANIFEST.json.

### Computational Environment

Heavy compute tasks (notebook execution, figure regeneration, and statistical testing) were performed on a Google Colab Pro environment with an NVIDIA A100 GPU. The local machine (MacBook Air M2, 8 GB RAM) functioned solely as a client for the web browser, file management (via GitHub and Google Drive), PDF rendering, and visual inspection. No resource-intensive operations were executed locally.

## 3. Results

We evaluated 17 eligible language models on molecular inputs across two primary domains: liver tissue transcriptomics and plasma cell-free RNA (cfRNA). The overarching conclusion is that under strictly evaluated, held-out conditions, language models do not demonstrate a reliable capability to predict MASLD fibrosis stages from molecular panels. Three primary findings emerge from our evaluation. First, none of the eligible tissue models exceeded the majority-class baseline on common binary items, and model rankings were highly unstable across different answer formats. Second, while some models achieved low Mean Absolute Error (MAE) in regression tasks, this was an artifact of reduced prediction variability (models regressing to the mean) rather than actual biological learning. Third, in cfRNA tasks, while some classical baselines narrowly passed our confidence-interval criteria, no language model demonstrated a statistically significant paired superiority over a constant majority classifier. Finally, our audit reveals that apparent model performance is heavily confounded by train-test data leakage in fine-tuned models and output truncation caused by evaluation harness limits.

### 3.1. Corpus composition

We aggregated the in-scope prompt collection from the source families (Table 1; Figure 1). The collection contains 106,688 in-scope prompts across 19 datasets; the full manifest carries 22 datasets and 107,004 items, of which 316 belong to datasets reserved for separate studies. The large count is dominated by auxiliary material: of the 92,721 training prompts, 90,533 are auxiliary knowledge, chemistry, geometry and training-side datasets, and 2,188 belong to the staged-liver and DRUG-seq pools; the remaining 13,967 prompts are frozen test prompts, including 504 geometric test prompts, and the pairwise formats expand a fixed set of profiles combinatorially.

**Figure 1.**
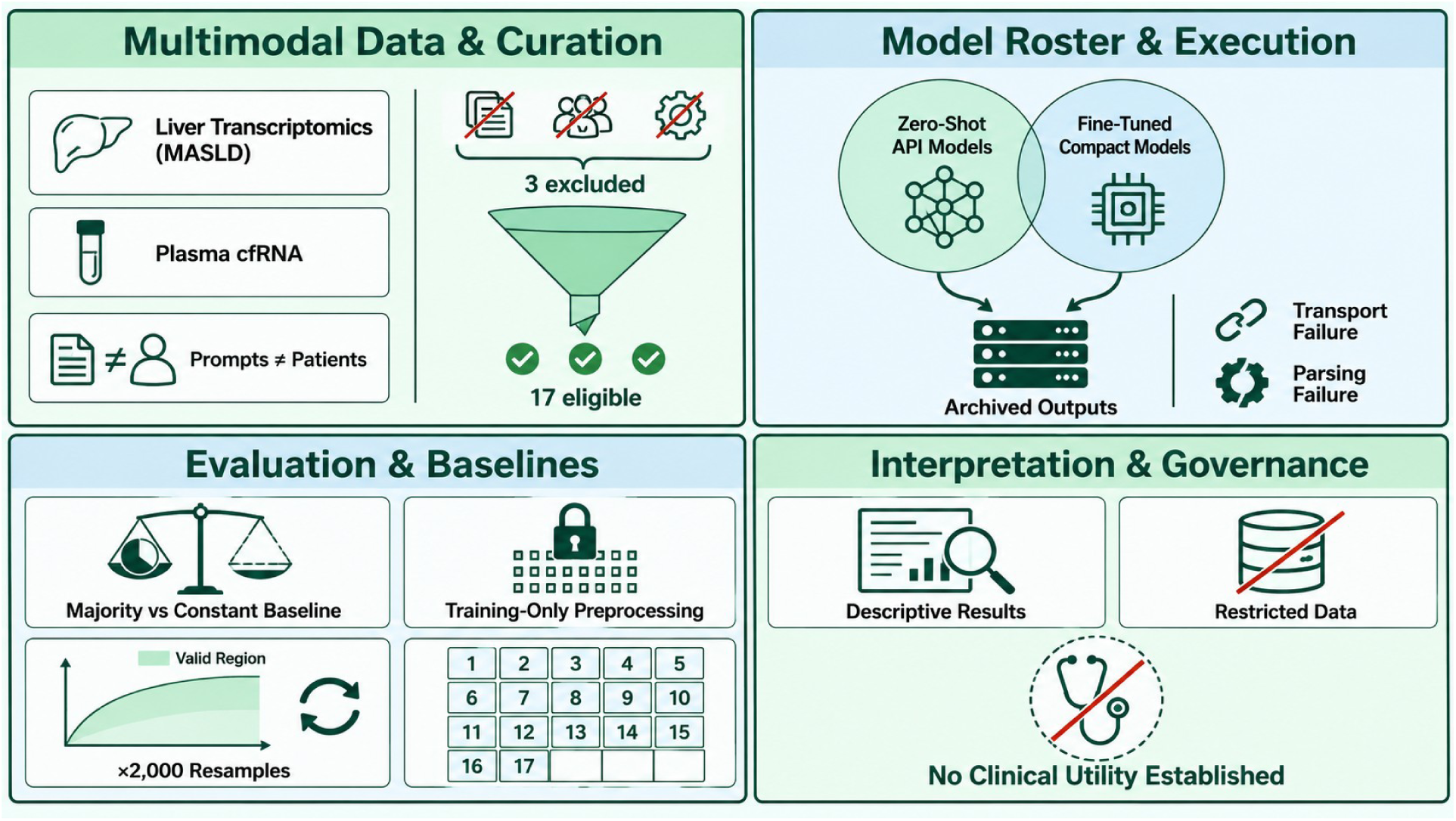
Data preparation, model evaluation and interpretation. The 17 eligible models and three excluded model versions refer to the tissue analysis; the excluded versions were fine-tuned on tissue test samples. Prompt counts are not patient counts. Classification is compared with majority-class baselines, and regression with training-fold median baselines. Training-only preprocessing is evaluated alongside full-dataset preprocessing. Intervals use 2,000 bootstrap resamples where specified. The analysis is retrospective and does not establish clinical utility. GPT-6 Astra (OpenAI) was used to assist in the conceptualization and drafting of this workflow schematic.

**Table 1.** Composition of the MASLD-Bench prompt collection by data source, domain and role. Counts from prompts/MANIFEST.json (v3-freeze-17).

| **Source** | **Domain** | **Train prompts** | **Test prompts** | **Formats** | **Role** |
|---|---|---|---|---|---|
| EPoS liver biopsies (GSE135251) | liver transcriptomics | — | 11,653 | binary, 5-class, ordinal, pairwise | frozen test |
| GSE135251 / GSE167523 / GSE130970 | staged liver | 1,196 | — | 5-class, ordinal, 3-class | SFT pool (shares GSE135251 samples with the frozen tissue test; see Methods) |
| GSE185051 | progression proxy | — | 104 | binary, ordinal | frozen test |
| GSE267032 | whole blood | — | 57 | 3-class, binary, ordinal | frozen test |
| MSV000094959 | liver→plasma proteome | — | 422 | 3-class, masked regression | frozen test |
| GSE247407 | primary hepatocyte | — | 30 | binary (FFA) | frozen test |
| E-MTAB-15601 | HepG2 DRUG-seq | 992 | — | 40-class, binary | SFT pool |
| S-BSST1614 cohort 1 (UZA + AMC) | plasma cfRNA | — | 660 | binary, 5-class, ordinal | frozen test + reader control |
| S-BSST1614 cohort 2 | plasma cfRNA | — | 537 | binary, 5-class, ordinal | frozen test |
| MB-40xx geometric test | expression geometry | — | 504 | geometric | frozen test (no exposed-checkpoint archive) |
| MB-10xx/11xx/ 61xx–63xx/70xx/ 8xxx, MB-40xx train | knowledge, chemistry, geometry | 90,533 | — | knowledge, drug–target, geometric | auxiliary |

The 106,688-prompt count (13,967 test and 92,721 training prompts) is a count of generated prompts, not a sample-size claim. The evaluated subset is 15–414 archived items per model–task combination, drawn from 21– 179 biological samples per family. Derived prompts are not independent biological observations, and we report patients and donors separately from prompt counts wherever a claim depends on sample size.

Corpus preparation included corrections to label parsing and a blood-proxy header alignment, and rejection of a cohort whose sample keys could not be verified. The current analysis uses the fixed, versioned prompt datasets and explicit archive-eligibility rules. The historical correction log is retained separately; incomplete documentation of source files and processing steps is disclosed in Section 5.

### 3.2. Eligible tissue models do not exceed majority-class accuracy

The primary tissue analysis uses an explicitly eligible model set: every model with all five tissue tasks, minus the three checkpoints whose SFT mixture contained the GSE135251 test samples. This leaves 17 models. For each single-sample task, scores are computed on the items present in every eligible archive (163 for binary and five-class, 162 for fibrosis regression and 163 for NAS regression). Invalid answers count as errors for classification. Regression MAE is conditional on a parsed number; valid-answer coverage, in-domain counts and a failure-aware sensitivity analysis are reported in Supplementary Table S2. Pairwise responses were not collected on one universal item set, so pairwise results are reported within their exact recorded evaluation sets and are not included in the cross-format mean rank.

On the 163 common binary items, the majority rate is 107/163 = 0.656. Six eligible models return exactly 0.656 and none exceeds it (Figure 3A). On the 163 common five-class items, Nemotron 120B is highest at 0.276 (95% CI 0.215–0.344), compared with a majority-class accuracy of 43/163 = 0.264 and uniform-choice chance accuracy of 0.200. The interval includes the majority rate; this marginal comparison is not a paired superiority test or evidence of equivalence. On the shared 300-item pairwise evaluation set used by 14 eligible models, Claude Sonnet 4.5 reaches 0.610 (sample-cluster bootstrap 95% CI 0.503–0.708) and Qwen3-0.6B v3 reaches 0.593 (0.491–0.693). DeepSeek and Nemotron share a different 258-item evaluation set within the dataset (0.419 and 0.500); Gemini has a separate 414-item evaluation set with almost no valid answers. These pairwise values are descriptive within their respective evaluation sets, not a single cross-model leaderboard.

Exposed-checkpoint diagnostic (not generalization evidence). On their own complete 168-sample and shared 300-pair evaluation sets, Qwen3-0.6B v6 scores 0.661 / 0.190 / 0.533 / 1.292 / 3.324, Qwen3-1.7B v6 scores 0.667 / 0.167 / 0.510 / 1.226 / 6.1×10¹², and Qwen3-8B v7 scores 0.661 / 0.220 / 0.483 / 1.205 / 1.547 across binary accuracy, five-class accuracy, pairwise accuracy, fibrosis MAE and NAS MAE. These values are retained as an audit of the trained models’ outputs on samples seen in SFT; they do not enter Tables 2A and 2B’s rankings or the Abstract’s claims. These exposed-checkpoint scores are descriptive diagnostics only.

**Table 2A.** Zero-shot API-evaluated models. Accuracy for MB-2001, MB-2002 and MB-2003; MAE among parsed numerical responses for MB-2005 and MB-2006. Mean ranks retain the original ranking across all 17 eligible models and are not recalculated within groups. The three exposed checkpoints are excluded and reported separately below. Pairwise cells show the number of evaluated items; values from different evaluation sets are not ranked together. The mean rank is over the four single-sample tasks only, with no-valid-numeric-output assigned the worst rank. MAE values above 100 indicate numeric-format failure, not ordinal error. Per-model valid/n and coverage of predictions within the permitted score range are in Supplementary Table S2.

| **Model** | **MB-2001 acc (n=163)** | **MB-2002 acc (n=163)** | **MB-2003 acc (evaluated n)** | **MB-2005 MAE (n=162)** | **MB-2006 MAE (n=163)** | **Mean rank** |
|---|---|---|---|---|---|---|
| GPT-4.1-mini | 0.656 | 0.190 | 0.370 (300) | 1.790 | 4.215 | 5.25 |
| Qwen2.5 72B | 0.656 | 0.166 | 0.253 (300) | 1.617 | 4.067 | 6.25 |
| DeepSeek Chat | 0.632 | 0.172 | 0.419 (258) | 1.778 | 3.994 | 6.75 |
| GPT-4.1 | 0.571 | 0.190 | 0.277 (300) | 1.802 | 4.061 | 7.00 |
| Nemotron 120B | 0.656 | 0.276 | 0.500 (258) | 2.074 | 3.833 | 3.75 |
| GPT-4o-mini | 0.509 | 0.178 | 0.187 (300) | 2732.179 | 4.111 | 9.50 |
| Mercury-2.5 | 0.583 | 0.172 | 0.437 (300) | 3082.712 | 5.981 | 10.00 |
| Qwen3 235B | 0.454 | 0.184 | 0.040 (300) | 14.210 | 28.374 | 10.00 |
| Llama 3.3 70B | 0.190 | 0.147 | 0.153 (300) | 1.517 | 2462.032 | 11.25 |
| Claude Sonnet 4.5 | 0.049 | 0.006 | 0.610 (300) | 27659.182 | 3.828 | 12.50 |
| Nova Pro | 0.166 | 0.172 | 0.330 (300) | 6789.636 | 9480.441 | 13.00 |
| GPT-5.6 Luna | 0.067 | 0.000 | 0.570 (300) | 1.800 | — | 14.50 |
| Gemini 2.5 Pro | 0.368 | 0.153 | 0.005 (414) | — | — | 15.25 |

**Table 2B.** Fine-tuned compact models. Metrics and mean ranks use the same definitions and overall 17-model ranking as Table 2A.

| **Model** | **MB-2001 acc (n=163)** | **MB-2002 acc (n=163)** | **MB-2003 acc (evaluated n)** | **MB-2005 MAE (n=162)** | **MB-2006 MAE (n=163)** | **Mean rank** |
|---|---|---|---|---|---|---|
| Qwen3-1.7B v3 | 0.656 | 0.190 | 0.470 (300) | 1.426 | 2.221 | 2.25 |
| Qwen3-1.7B v5 | 0.656 | 0.245 | 0.510 (300) | 1.142 | 6.1×10^26 | 4.50 |
| Qwen3-0.6B v3 | 0.399 | 0.202 | 0.593 (300) | 1.481 | 6.1×10^26 | 8.00 |
| Qwen3-0.6B v5 | 0.656 | 0.196 | 0.467 (300) | 2.1×10^12 | 213.086 | 8.00 |

**Table 3.** Binary class distributions and baselines for the plasma cfRNA tasks.

| **Task** | **Samples** | **Negative** | **Positive** | **Majority accuracy** |
|---|---|---|---|---|
| MB-8901 reference cohort (UZA) | 112 | 80 | 32 | **0.7143** |
| MB-8911 second cohort | 179 | 95 | 84 | **0.5307** |
| MB-8921 AMC variant | 108 | 73 | 35 | **0.6759** |

**Table 4.**
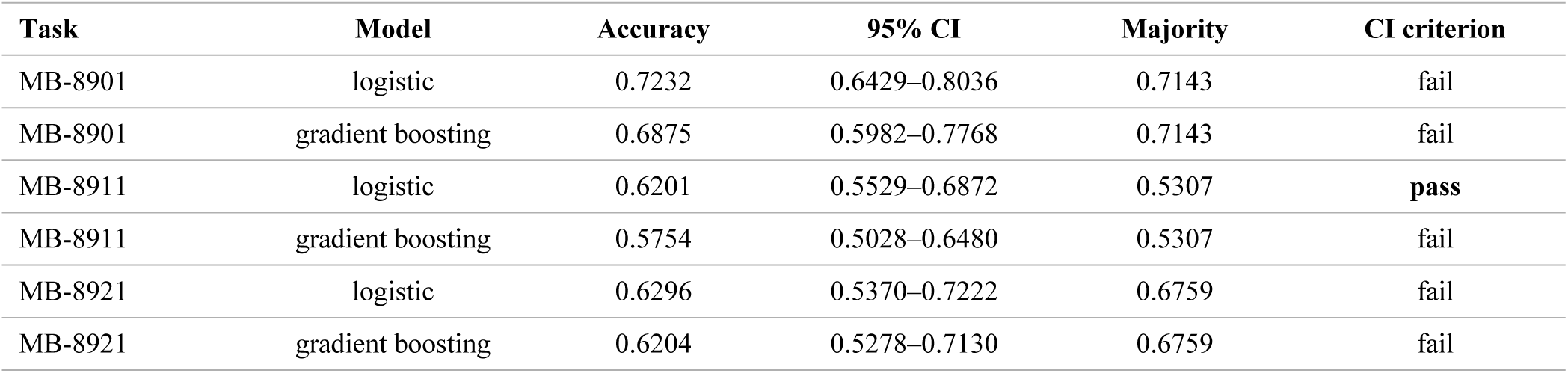
Classical confidence-interval criterion decisions.

| Task | Model | Accuracy | 95% CI | Majority | CI criterion |
|---|---|---|---|---|---|
| MB-8901 | logistic | 0.7232 | 0.6429–0.8036 | 0.7143 | fail |
| MB-8901 | gradient boosting | 0.6875 | 0.5982–0.7768 | 0.7143 | fail |
| MB-8911 | logistic | 0.6201 | 0.5529–0.6872 | 0.5307 | **pass** |
| MB-8911 | gradient boosting | 0.5754 | 0.5028–0.6480 | 0.5307 | fail |
| MB-8921 | logistic | 0.6296 | 0.5370–0.7222 | 0.6759 | fail |
| MB-8921 | gradient boosting | 0.6204 | 0.5278–0.7130 | 0.6759 | fail |

**Table 5.**
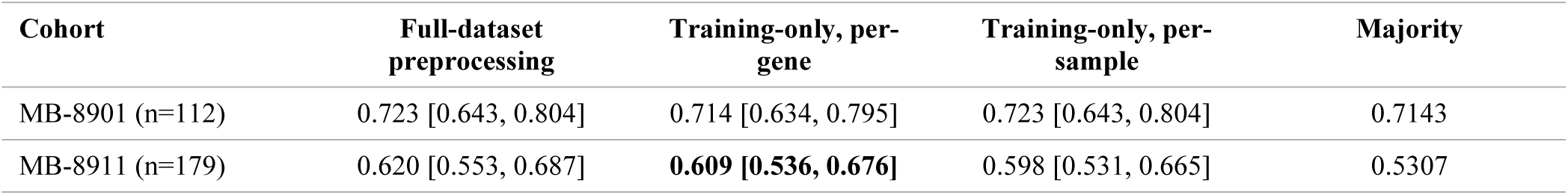
Representation sensitivity of the binary classical result.

**Table 6.** Best language-model result per binary cfRNA task against the corrected majority rate.

| **Task** | **Best model** | **Accuracy [95% CI]** | **Valid/n** | **Sensitivity** | **Specificity** | **Majority** |
|---|---|---|---|---|---|---|
| MB-8901 reference cohort | GPT-4o-mini | 0.670 [0.580, 0.759] | 105/112 | 0.000 | 0.938 | 0.714 |
| MB-8911 second cohort | Qwen3-235B | 0.592 [0.520, 0.665] | 179/179 | 0.726 | 0.474 | 0.531 |

**Table 7.** Complete confusion counts, separating valid answers from non-answers. The valid-only matrix uses answered items only. Invalid answers and API request failures are then counted by reference class, so a reader can reconstruct either a valid-only or a failure-inclusive view without recomputing anything.

| **Model** | **Task** | **TP** | **FN** | **TN** | **FP** | **Invalid on positive** | **Invalid on negative** | **Transport error on positive** | **Transport error on negative** |
|---|---|---|---|---|---|---|---|---|---|
| GPT-4o-mini | 8901 | 0 | 30 | 75 | 0 | 2 | 5 | 0 | 0 |
| GPT-4o-mini | 8911 | 0 | 83 | 93 | 0 | 1 | 2 | 0 | 0 |
| GPT-4.1-mini | 8901 | 3 | 29 | 68 | 12 | 0 | 0 | 0 | 0 |
| GPT-4.1-mini | 8911 | 5 | 79 | 92 | 1 | 0 | 2 | 0 | 0 |
| Qwen3-235B | 8901 | 26 | 6 | 26 | 54 | 0 | 0 | 0 | 0 |
| Qwen3-235B | 8911 | 61 | 23 | 45 | 50 | 0 | 0 | 0 | 0 |
| Nemotron 120B | 8901 | 0 | 0 | 0 | 0 | 0 | 0 | 32 | 80 |
| Nemotron 120B | 8911 | 0 | 0 | 0 | 0 | 0 | 0 | 84 | 95 |

**Table 8.** Ordinal baselines with the original estimators and the original AMC gene-selection and standardisation procedure. The estimators are Ridge(alpha=1.0) and HistGradientBoostingRegressor. Both rounded stage predictions and continuous predictions are reported. For AMC, the primary procedure constructs the panel and per-gene means and standard deviations on all 112 reference cohort samples before subsetting to the 108 jointly staged samples. † For MB-8925 the panel and per-gene means and standard deviations are estimated on all 112 reference cohort samples and the resulting feature matrix is then subset to the 108 AMC-staged rows, exactly as scripts/gates_mb89xx.py does (Xa = X1[amc_mask]). A sensitivity analysis that rebuilds the panel on the 108 samples alone is reported in §3.7 and gives the same qualitative result.

| Task | Estimator | Prediction | MAE | Constant MAE | Paired difference [95% CI] | Improvement demonstrated? |
|---|---|---|---|---|---|---|
| MB-8905 | ridge | rounded | 1.134 | 1.250 | +0.116 [-0.085, +0.326] | no |
| MB-8905 | ridge | continuous | 1.255 | 1.250 | -0.005 [-0.206, +0.189] | no |
| MB-8905 | hgb regressor | rounded | 1.152 | 1.250 | +0.098 [-0.036, +0.241] | no |
| MB-8905 | hgb regressor | continuous | 1.144 | 1.250 | +0.106 [-0.027, +0.237] | no |
| MB-8915 | ridge | rounded | 1.626 | 1.341 | -0.285 [-0.503, -0.061] | no |
| MB-8915 | ridge | continuous | 2.121 | 1.341 | -0.780 [-1.028, -0.523] | no |
| MB-8915 | hgb regressor | rounded | 1.240 | 1.341 | +0.101 [-0.017, +0.212] | no |
| MB-8915 | hgb regressor | continuous | 1.253 | 1.341 | +0.087 [-0.023, +0.195] | no |
| MB-8925† | ridge | rounded | 1.148 | 1.019 | -0.130 [-0.315, +0.074] | no |
| MB-8925† | ridge | continuous | 1.226 | 1.019 | -0.208 [-0.380, -0.024] | no |
| MB-8925† | hgb regressor | rounded | 0.963 | 1.019 | +0.056 [-0.065, +0.176] | no |
| MB-8925† | hgb regressor | continuous | 0.952 | 1.019 | +0.067 [-0.049, +0.180] | no |

### 3.3. Model rankings differ across answer formats

Figures 2, 4 and S5 display the rank patterns and their spread. Model rankings were not stable across answer formats applied to the same frozen samples (Figure 4). Within the eligible set and on the common item sets, Claude Sonnet 4.5 moves from seventeenth of 17 on binary fibrosis (0.049) to first of 14 on the shared pairwise evaluation set (0.610); Qwen3-0.6B v3 moves from twelfth on binary (0.399) to second on pairwise (0.593). GPT-5.6 Luna is third on pairwise (0.570) but sixteenth on binary (0.067) and last on five-class (0.000). The pattern is not evidence that format causes a capability change: the formats ask for different computations on the same measurements — a threshold, a five-category assignment, a comparison and a numerical value — and the models differ in training exposure.

**Figure 2.**
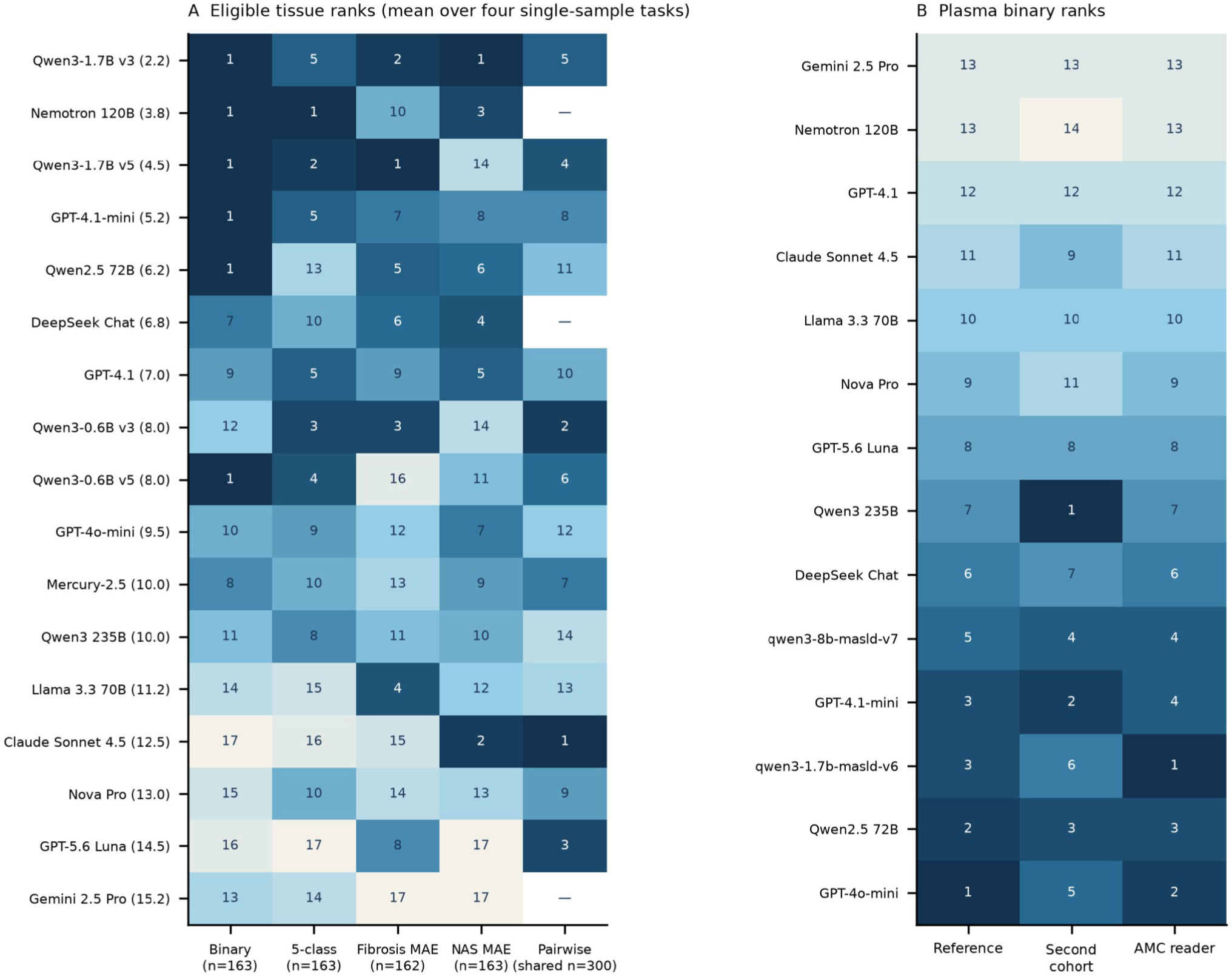
Model ranks on common evaluated items. Panel A shows 17 eligible tissue models across four single-sample tasks and pairwise ranks only for models sharing the 300-item evaluation set. Mean ranks exclude pairwise tasks. Panel B shows cfRNA binary ranks, with AMC scores obtained by rescoring fixed predictions on 108 samples. Ranks do not imply superiority over majority baselines; evaluations with API request failures do not measure capability.

**Figure 3.**
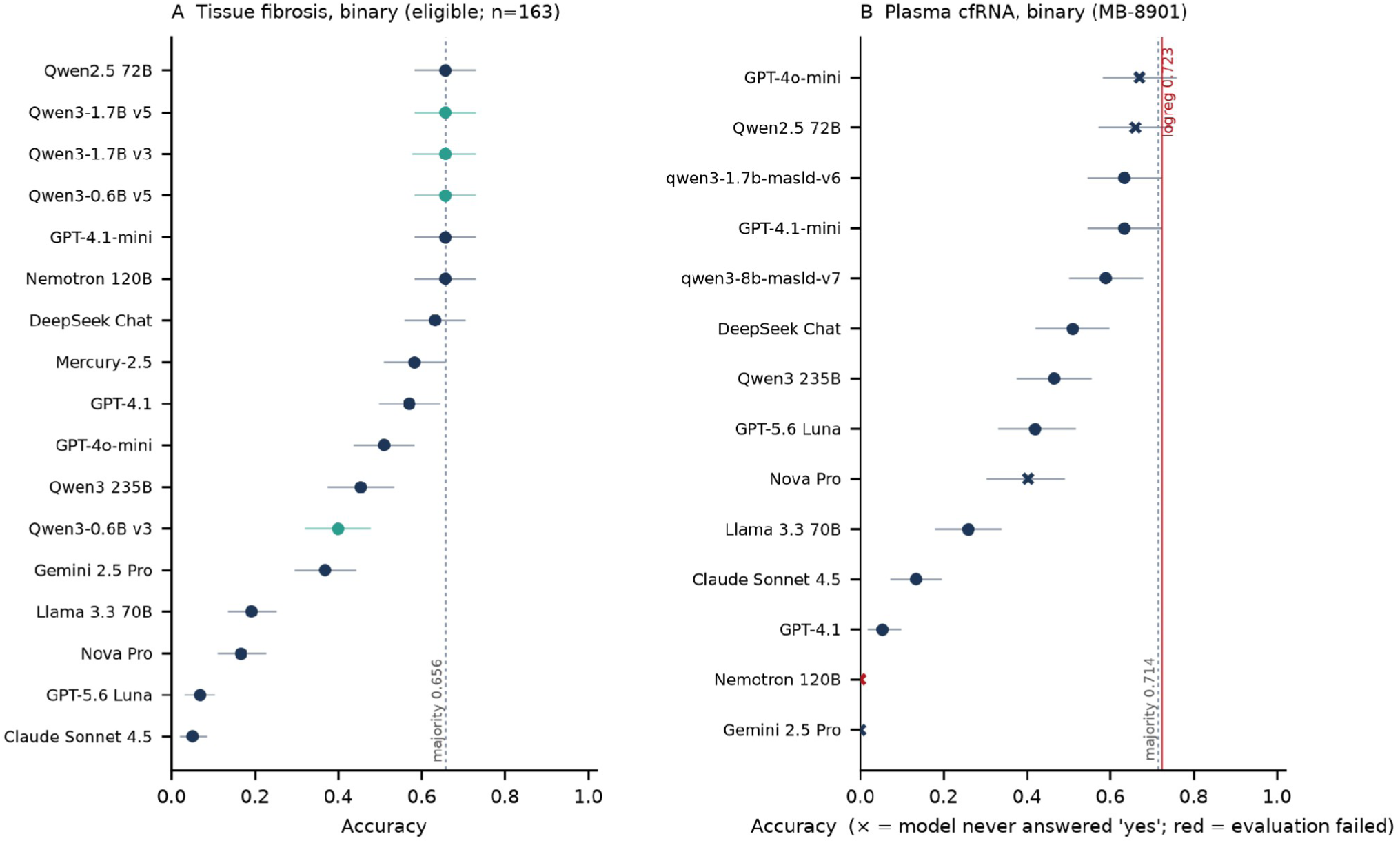
Binary accuracy and intervals. Panel A uses the 17 eligible tissue models on 163 aligned items and observed majority 107/163 = 0.656. Panel B uses the cfRNA reference cohort (n = 112), with majority 0.714 and frozen-representation logistic accuracy 0.723. Crosses indicate no positive answer; red markers identify failed evaluations. Intervals and point estimates do not establish paired superiority.

**Figure 4.**
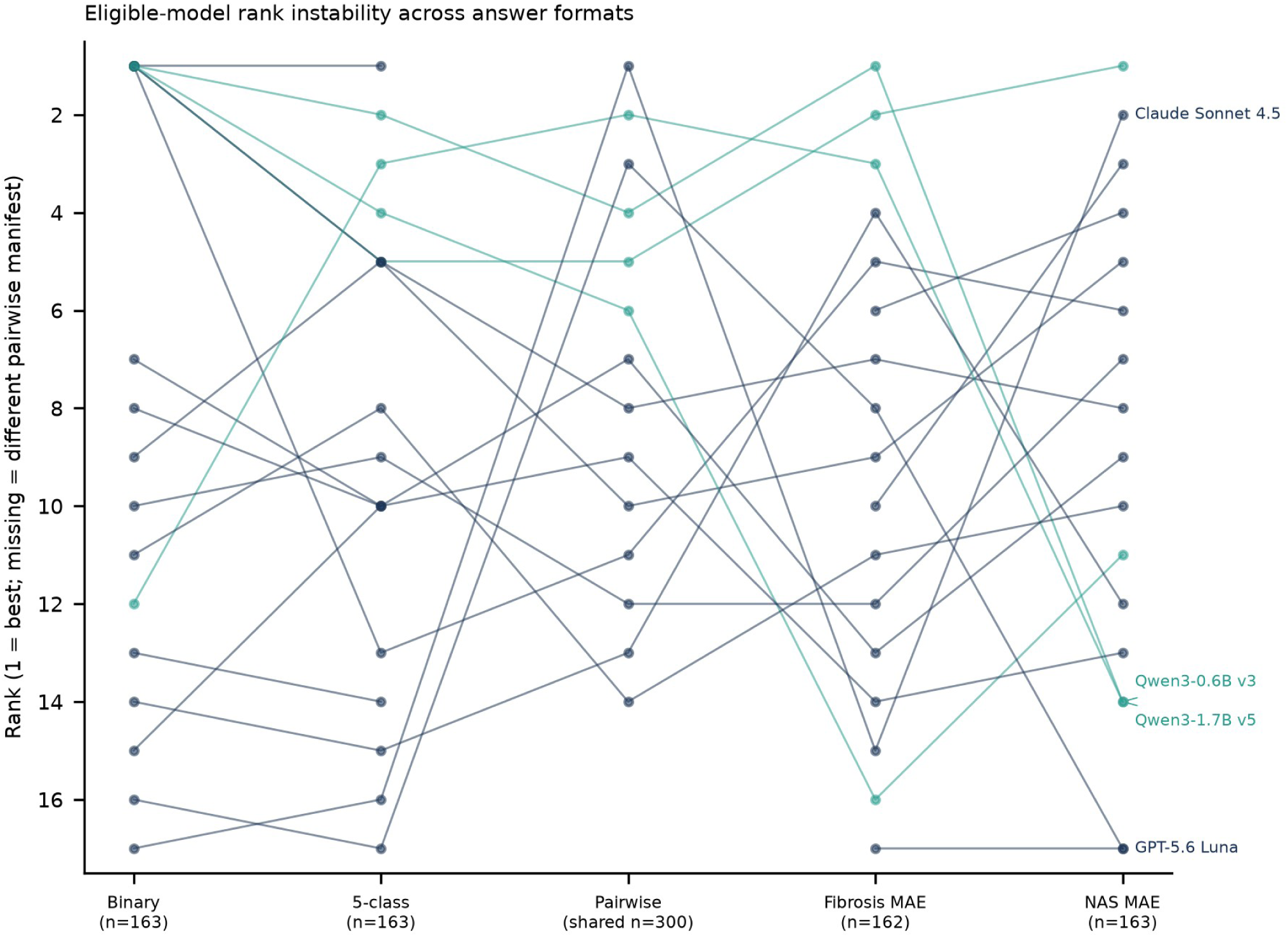
Rank variation across answer formats in the eligible tissue model set. Single-sample ranks use aligned items; pairwise ranks use only the shared 300-item evaluation set. Teal denotes compact v3/v5 models and navy API systems. Missing pairwise ranks indicate a different evaluation set. This is a descriptive comparison, not a causal effect of format or scale.

The eligible comparison still crosses compact and API-accessed model classes with different training histories, and pairwise ranks are computed only within a shared recorded evaluation set. What the data support is narrower: within this model set, changing the answer format reorders models substantially, and aggregate leaderboards conceal that instability. No causal claim about format or model scale is made.

The eligible five-class distribution is shown in Figure S2. A developer-level comparison (Figure S1) is descriptive only and must be read with the same eligible-model and denominator restrictions. It should not be used to claim that fine-tunes as a class dominate API-accessed models: the exposed v6/v7 checkpoints are excluded from the primary rankings, and numeric-format failures remain visible rather than being converted into evidence of staging skill.

### Fibrosis Stage Classification Tasks

MB-2001 is binary advanced fibrosis (F3–F4 versus F0–F2). MB-2002 is five-stage classification (F0, F1, F2, F3, F4). MB-2003 is pairwise stage comparison between two profiles. MB-2005 is numeric prediction of fibrosis stage. MB-2006 is numeric prediction of the NAFLD Activity Score (NAS). NAS is a composite histological activity score; both MB-2005 and MB-2006 are ordinal scales treated as numeric targets, which is a modelling convenience rather than a claim that the underlying quantity is continuous.

### 3.4. Why Low Error Rates Are Misleading in Regression Tasks?

Figures 5 and S6 show error/correlation and distributions of predictions within the permitted score range; Figure S7 shows valid-answer coverage.

**Figure 5.**
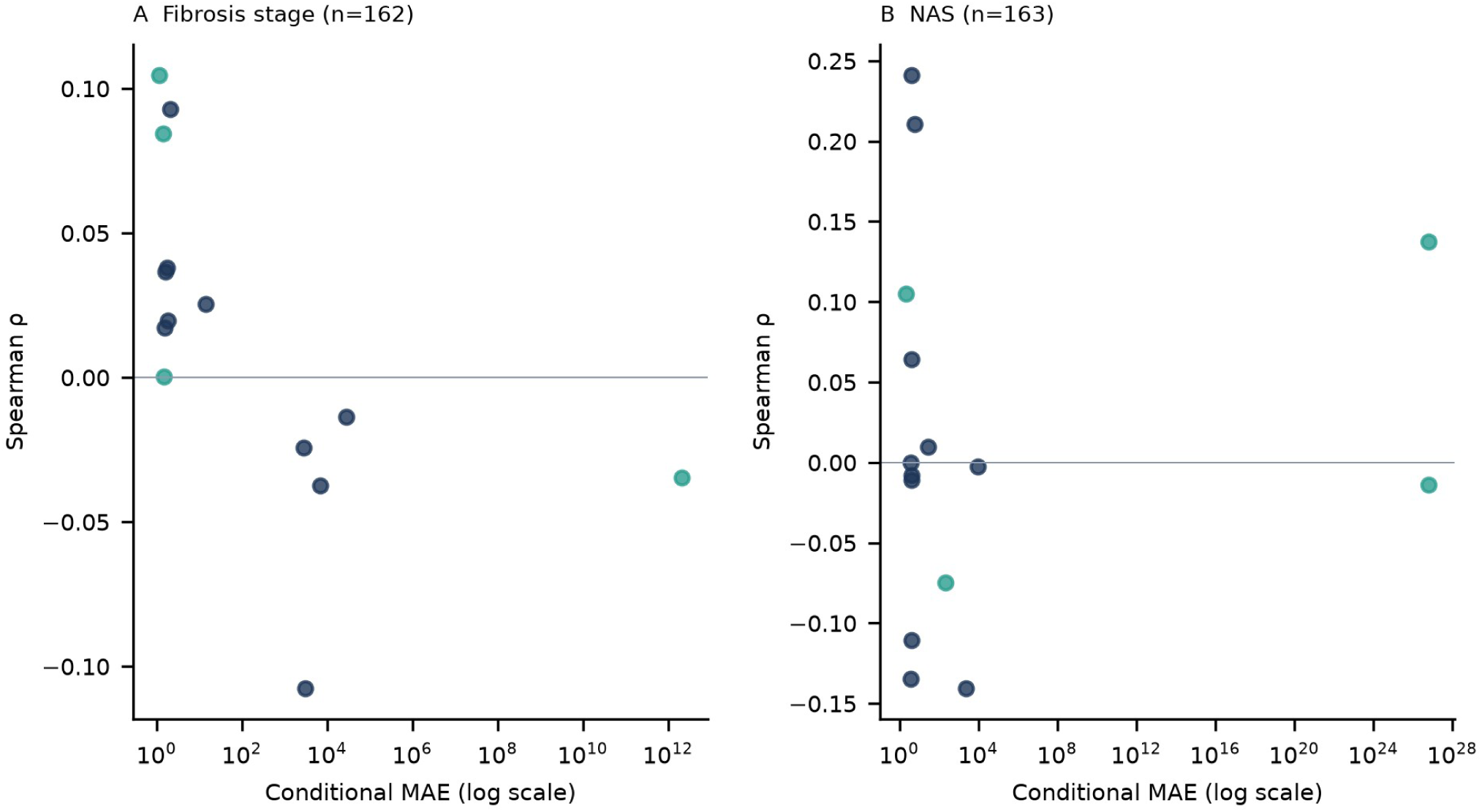
MAE among parsed numerical responses on a logarithmic axis versus Spearman correlation for eligible tissue models. Fibrosis uses 162 common archived items and NAS 163; parsed denominators differ by model. Teal denotes compact models and navy API systems. Extreme MAE reflects numeric-format failures; absent numeric predictions are not plotted. Supplementary Table S2 provides model-specific values and coverage.

Within the 162-item fibrosis-regression set archived for every eligible model, the lowest eligible compact conditional MAE is Qwen3-1.7B v5 at 1.142 (162/162 parsed), and the lowest API-model conditional MAE is Llama 3.3 70B at 1.517 (150/162 parsed). Their Spearman correlations are 0.105 and 0.017 respectively. Because the conditional denominators differ, the direct analysis restricted to items parsed for both models uses the 150 items parsed by both models: MAE 1.127 for Qwen3-1.7B v5 and 1.517 for Llama. This subset is selected by successful parsing and does not replace failure-aware whole-set reporting. The comparison is descriptive, not a test of compact-versus-API-model superiority. On NAS regression, Qwen3-1.7B v3 has MAE 2.221 with ρ = 0.105, while several models fail numerically.

The error and the information content must be separated. For Qwen3-1.7B v5 on MB-2005, the observed mean is 1.802 (SD 1.230) and the predicted mean is 1.315 (SD 0.664); for Llama, the predicted mean is 0.477 (SD 0.519). Predictions are compressed relative to the observed distribution, but they do not simply equal the cohort mean. We therefore describe the pattern as reduced prediction variability with weak rank correlation: an error metric can improve while the model recovers little biological ordering. The mechanism remains unresolved. It means the model is essentially ’hedging its bets’ by guessing the cohort average, rather than actually distinguishing between a healthy liver and a fibrotic one.

Conditional MAE is not a complete evaluation of these outputs. We therefore also report coverage of predictions within the permitted score range and an MAE with penalties for missing or out-of-range predictions in Supplementary Table S2. Values outside the declared domains (0–4 for fibrosis and 0–8 for NAS) are not treated as plausible staging estimates. Qwen3-1.7B v5, for example, produces extreme numeric failures on NAS even though its fibrosis MAE is the lowest eligible value. No single MAE is presented as evidence of clinical or biological staging competence.

### 3.5. Binary cfRNA baselines and preprocessing sensitivity

Figures 3 and S3 show binary performance; Figure S4 summarizes the cfRNA cohort and preprocessing procedures.

The binary tasks are MB-8901 (UZA reference cohort, 112 samples), MB-8911 (second cfRNA cohort, 179 samples) and MB-8921 (AMC-reader variant, 108 samples).

The binary majority rate is computed from the evaluated binary labels, independently of the five-class distribution.

In the reference cohort, logistic regression achieves accuracy of 0.723 (95% CI 0.643–0.804); the lower confidence limit does not exceed the majority-class accuracy of 0.714. Histogram gradient boosting achieves 0.688 and also does not meet this criterion. Neither model meets the criterion using the AMC reference labels (accuracies 0.630 and 0.620 versus majority-class accuracy 0.676). In the second cfRNA cohort, logistic regression achieves 0.620 (95% CI 0.553–0.687), with **a lower** confidence limit above 0.531. Gradient boosting achieves 0.575 (95% CI 0.503–0.648) and does not meet the criterion.

Under the procedure using full-dataset preprocessing, second cfRNA cohort logistic accuracy is 0.620 [0.553, 0.687]. With training-only panel selection and per-gene means and standard deviations it is 0.609 [0.536, 0.676]; with training-only panel selection and per-sample standardisation it is 0.598 [0.531, 0.665]. The gene-wise lower bound exceeds the observed majority by approximately 0.5 percentage points, whereas the per-sample lower bound equals it at full precision. These threshold outcomes do not establish a difference between pipelines.

Three preprocessing procedures are distinguished. Full-dataset preprocessing selects the top-200 gene panel and calculates per-gene means and standard deviations using all samples, including held-out samples. Training-only per-gene preprocessing selects the panel and estimates these quantities using only the training samples. Training-only per-sample preprocessing selects the panel using training samples and standardises values across the selected genes within each sample. All three procedures use the same five folds and seed.

In the reference cohort, none of the three preprocessing procedures produces an accuracy confidence interval whose lower limit exceeds the majority-class accuracy. In the second cfRNA cohort, this criterion is met with full-dataset preprocessing and training-only per-gene preprocessing, but not with per-sample standardisation. No paired comparison between preprocessing procedures was performed.

### 3.6. Selected language-model comparisons do not establish a majority-classifier advantage

We re-parsed every archived cfRNA response with the same parser used for the original scoring, and separated API request failures from invalid model answers from valid but incorrect predictions.

On MB-8901, GPT-4o-mini returns 105 negative answers and seven unparseable responses, with no positive answers. The valid-answer confusion counts are TP = 0, FN = 30, TN = 75 and FP = 0; two additional positive and five negative items are unparsed. Failure-inclusive sensitivity is 0/32 = 0.000, specificity is 75/80 = 0.938 and balanced accuracy is 0.469. Accuracy is 75/112 = 0.670 and does not indicate detection of advanced fibrosis.

Qwen2.5 72B shows the same reference cohort pattern (0.661, sensitivity 0.000). On the second cfRNA cohort, two models sit above the majority rate numerically: Qwen3-235B at 0.592 and GPT-4.1-mini at 0.542 against 0.531. Numerical superiority is not statistical superiority. For Qwen3-235B the paired comparison against always answering "no" gives 61 wins and 50 losses, exact McNemar p = 0.3426. Counting invalid answers as model errors, the reference cohort comparison gives 0 wins and 5 losses (p = 0.0625); Holm-adjusted across these two comparisons the values are 0.125 and 0.343. Neither is significant, and both models were selected as the best performers on the same sets, so the tests are post-selection and exploratory. The declared family covers two selected comparisons, not the full set of 14 models, and no family-wise claim across the model set is made. These selected exploratory comparisons do not establish a language-model advantage over the majority classifier; they do not prove that every model lacks all discrimination.

GPT-4o-mini’s reference cohort row is the decisive one: 0 true positives, 30 false negatives, 75 true negatives, 0 false positives, plus 2 unparseable positives and 5 unparseable negatives. Its 0.670 accuracy is 75/112.

### Validity varies enough to matter

Across the 14 cfRNA archives, parsed-answer coverage on the reference cohort ranges from 0/112 to 112/112. Gemini 2.5 Pro returned parseable answers for 2 of 112 items; GPT-4.1 for 14 of 112; Nova Pro for 59. Those accuracies are reported as observed but are not interpretable as capability, because a model that rarely produces an answer is being scored mostly on its failures. The evaluation script’s 24-token compact budget and 900-token API-model budget contribute to truncation and are stated in Section 2.2.

### Operational failure

All 873 cfRNA archive rows for Nemotron 120B contain a API request failure and no valid completion. These rows describe a failed evaluation and are excluded from capability comparisons; they do not characterize the model’s tissue-task performance.

### 3.7. Fibrosis-stage prediction compared with a training-fold median baseline

The MAE of each ordinal model is compared with that of a baseline that predicts the training-fold median for every held-out sample. The comparison uses the baseline prediction error, not the numerical value of the median stage.

No evaluated ordinal estimator demonstrates lower MAE than the training-fold median baseline under the specified paired-interval criterion. On MB-8905, rounded Ridge MAE is 1.134 versus 1.250, with paired difference +0.116 [−0.085, +0.326]. On MB-8915, rounded Ridge is worse than the constant (1.626 versus 1.341), while rounded gradient boosting has a positive but inconclusive difference (1.240 versus 1.341; +0.101 [−0.017, +0.212]). Under the original AMC procedure, rounded gradient boosting gives 0.963 versus 1.019 and Ridge gives 1.148; neither demonstrates an advantage. Rebuilding the representation on 108 AMC samples gives respective MAEs of 0.991 and 1.204, with the same qualitative conclusion.

### 3.8. Feature Selection and External Transfer Fail to Outperform Baselines

Figure 6 shows accuracy using different numbers of selected genes. Figure S8 distinguishes training-set accuracy, classifier cross-validation with a fixed panel, cross-validation that repeats all selection steps, and external evaluation.

**Figure 6.**
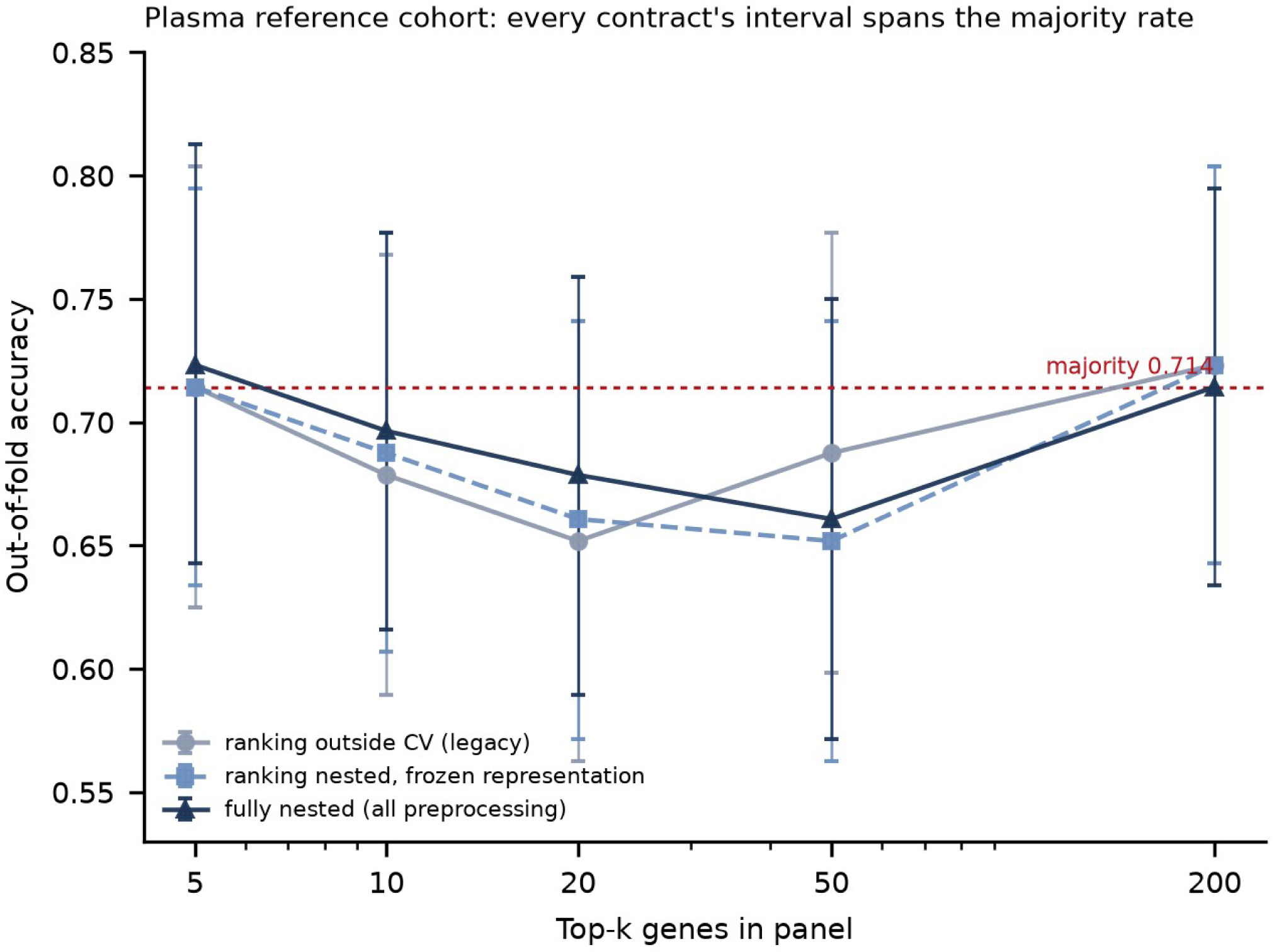
Top-k accuracy in the 112-sample cfRNA reference cohort. Grey uses ranking outside cross-validation; light blue nests ranking with full-dataset representation fixed; navy nests panel selection, ranking and scaling. Every 2,000-resample interval spans the majority rate 0.714. Intervals resample fixed out-of-fold predictions and do not include all model-fitting uncertainty.

We distinguish three top-k procedures: supervised ranking outside cross-validation; ranking nested within folds with a full-dataset panel and scaling; and fully nested panel selection, ranking and scaling. Only the third evaluates the complete feature-selection procedure without held-out samples entering preprocessing.

With supervised ranking moved inside the training fold but the full-dataset panel and scaling held fixed, the top-5 panel reaches 0.714 — exactly the majority rate of 80/112. Nesting the whole procedure, including panel selection and scaling, gives 0.723 at k = 5 and 0.714 at k = 200 (Table 9). The three procedures agree within a few items at every k, and every interval spans the majority rate (Table 9). These overlapping intervals do not establish equivalence between the procedures.

**Table 9.** Signal concentration under three preprocessing procedures, with bootstrap intervals. Each cell reports an accuracy and a 2,000-resample interval over fixed out-of-fold predictions. These intervals describe sampling variability of the recorded predictions, conditional on the realized folds and fits; they do not encompass all model-fitting or selection uncertainty.

| **Top-k genes** | **Ranking outside CV (legacy)** | **Ranking nested, frozen representation** | **Fully nested (all preprocessing)** |
|---|---|---|---|
| 5 | 0.714 [0.625, 0.804] | 0.714 [0.634, 0.795] | 0.723 [0.643, 0.813] |
| 10 | 0.679 [0.589, 0.768] | 0.688 [0.607, 0.777] | 0.696 [0.616, 0.777] |
| 20 | 0.652 [0.562, 0.741] | 0.661 [0.571, 0.741] | 0.679 [0.589, 0.759] |
| 50 | 0.688 [0.598, 0.777] | 0.652 [0.562, 0.741] | 0.661 [0.571, 0.750] |
| 200 | 0.723 [0.643, 0.804] | 0.723 [0.643, 0.804] | 0.714 [0.634, 0.795] |

The majority rate is 0.714. **Every interval in the table spans it**, including the fully nested top-5 point estimate of 0.723 [0.643, 0.813]. No procedure demonstrates a top-k advantage, under this confidence-interval criterion. For external transfer, the five genes selected in the reference cohort are aligned to the second cfRNA cohort matrix by Ensembl identifier. Column position is not a valid correspondence because the independently selected 200-gene panels share only nine genes.

The corrected analysis matches genes by Ensembl identifier, confirms that all five genes selected in the reference cohort are present in the second cohort, and applies the per-gene standardisation estimated in the reference cohort without modification. Accuracy on the reference-cohort samples used to fit the model is 0.759. External accuracy in the second cfRNA cohort is 0.615 (95% CI 0.547–0.687), compared with majority-class accuracy of 0.531; sensitivity is 0.607 (51/84) and specificity is 0.621 (59/95). Refitting the model using the same five genes within the second cohort gives cross-validated accuracy of 0.682.

The external paired comparison and the internal validation in the reference cohort estimate different quantities.

### The external comparison is descriptive and does not demonstrate superiority

A marginal accuracy interval that excludes a fixed observed majority rate is not the same procedure as a paired test against the majority classifier. On the same 179 second cfRNA cohort subjects the panel produces 51 wins and 36 losses against always-"no": exact McNemar p = 0.133, and the paired accuracy difference is +0.084 with a bootstrap interval of −0.017 to +0.184, which includes zero. The point estimate is above the majority rate and the marginal interval excludes it, but the paired comparison does not establish superiority at the 0.05 level. We therefore report "0.615 against 0.531" as a descriptive result and make no superiority claim.

### Internal cross-validation with a fixed panel does not validate panel selection

The reference-cohort accuracy of 0.714 comes from cross-validating only the classifier; the panel was selected once using all reference-cohort labels, and standardisation was calculated using all reference-cohort samples. Repeating gene selection by median absolute deviation, supervised ranking and standardisation within each training fold gives accuracy of 0.723 [0.643, 0.813]. Its lower confidence limit also does not exceed the majority rate of 0.714. Neither analysis establishes an accuracy advantage in the reference cohort. The first holds the selected panel fixed; the second evaluates a procedure that may select different genes in each fold.

The panel selected in the reference cohort has external accuracy descriptively above the second cfRNA cohort majority, without a demonstrated paired superiority. Neither the conditional fixed-panel analysis nor the fully nested procedure demonstrates a reference cohort advantage. Cohorts differ in label source and class balance; the five-gene panel remains exploratory and requires independent replication.

Single-gene descriptive AUROCs on the reference cohort are 0.716, 0.716, 0.698, 0.691 and 0.691 for the five strongest Ensembl features. Those are descriptive, not cross-validated, and they do not establish that any language model used those genes. No LLM feature-ablation experiment was run, and no mechanistic claim is made.

The cell-composition hypothesis — that plasma cfRNA partly reflects hepatocyte-derived RNA — remains a candidate explanation, not a tested mechanism. Cross-referencing the top-20 informative genes per task against the deposit’s cell-type Gini table found no strong markers in either corpus. Absence of overlap with a 152-gene marker list does not exclude a cell-composition explanation; it only fails to support one specific version of it.

### 3.9. Detection of advanced fibrosis and agreement between biopsy readers

#### Discrimination

Rescoring frozen ordinal outputs as advanced-fibrosis scores gives a model-level AUROC on the reference cohort of 0.628 for Qwen3-8B v7 (n = 112, 32 positive, average precision 0.366 against a prevalence of 0.286). On the second cfRNA cohort, rescoring restricted to predictions within the permitted range gives AUROC 0.641 for GPT-5.6 Luna (46/179 valid, 13 positive) and 0.581 for GPT-4.1-mini (106/179 valid, 49 positive). These values calculated only for responses with valid predictions are not a full-cohort comparison; the evaluated populations differ between models.

Two cautions apply. First, these are **discrimination** results on ordinal outputs, and they are not comparable with accuracy figures from a yes/no format. Second, coverage differs sharply between models, so an AUROC computed on 46 items is not the same quantity as one computed on 179.

Average precision is computed with a tie-aware implementation. It is not interchangeable with trapezoidal precision–recall area; score ties must not acquire an arbitrary ordering.

### Reader disagreement (Figure S9)

Two independent biopsy readings exist for 108 of the 112 reference cohort samples. Exact agreement is 56/108 (51.9%). Unweighted Cohen’s κ is 0.400; linearly weighted κ is 0.641; quadratic-weighted κ is 0.809; for the binary F3–F4 versus F0–F2 collapse, κ is 0.715. Agreement uses only the 108 samples staged by both readers.

A difference between reader-referenced accuracies is not a universal performance ceiling. Neither reader supplies adjudicated truth here, and the observed differences depend on model, endpoint and metric.

What can be said is narrower and is reported per model. With predictions held fixed on the same 108 samples, swapping the reference reader changes five-class accuracy for that one classical model from 0.389 to 0.259. For language models on the **binary** task, the change is much smaller: across all 14 archives, the largest difference between UZA-referenced and AMC-referenced accuracy is 4.6 percentage points (GPT-4.1-mini 0.630 versus 0.583), and most models move by less than one point (Table 10).

**Table 10.**
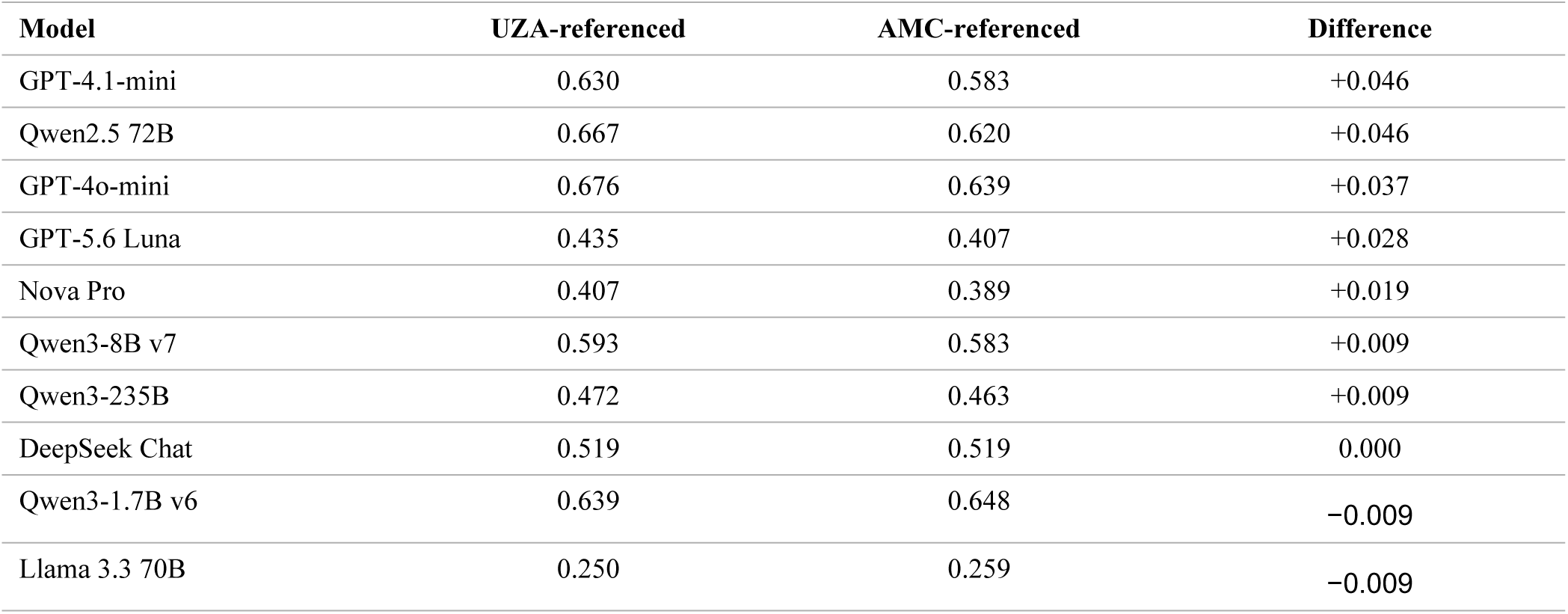
Per-model reader sensitivity on the 108 jointly staged samples (binary advanced fibrosis).

| Model | UZA-referenced | AMC-referenced | Difference |
|---|---|---|---|
| GPT-4.1-mini | 0.630 | 0.583 | +0.046 |
| Qwen2.5 72B | 0.667 | 0.620 | +0.046 |
| GPT-4o-mini | 0.676 | 0.639 | +0.037 |
| GPT-5.6 Luna | 0.435 | 0.407 | +0.028 |
| Nova Pro | 0.407 | 0.389 | +0.019 |
| Qwen3-8B v7 | 0.593 | 0.583 | +0.009 |
| Qwen3-235B | 0.472 | 0.463 | +0.009 |
| DeepSeek Chat | 0.519 | 0.519 | 0.000 |
| Qwen3-1.7B v6 | 0.639 | 0.648 | −0.009 |
| Llama 3.3 70B | 0.250 | 0.259 | −0.009 |

Reader disagreement is a real constraint on what any staging model can be shown to have learned. It is not a fixed ceiling, and its magnitude depends on the model, the endpoint and the metric.

### 3.10. Scope of additional experimental arms

Transfer, reasoning and retrieval arms were constructed but their archived numerical results were not re-derived in the authoritative analysis used here. They are outside the evidentiary scope of this report. No at-scale tool-orchestration evaluation is claimed.

## 4. Discussion

The primary tissue and cfRNA analyses do not establish reliable molecular staging by the evaluated language models under the archived protocols. Eligible tissue models do not exceed the binary majority accuracy on common items; low conditional regression error can coexist with weak rank correlation. A classical cfRNA result in the second cfRNA cohort meets the specified confidence-interval criterion under some preprocessing procedures, but this is not a language-model result or a clinical validation.

This evaluation demonstrates why model comparisons require an explicit baseline, an eligible model set and common item sets. A five-class majority is an inappropriate reference for binary accuracy, and a median stage label is not a constant predictor’s MAE. Training exposure is a different issue: three checkpoints include every tissue test sample in their fine-tuning pool and cannot supply held-out tissue evidence. These problems require artifact-level checks and cannot be resolved from an aggregate leaderboard alone.

Ranks differ across answer formats in these archived data: Claude Sonnet 4.5 ranks last on binary fibrosis and first within the shared pairwise evaluation set. Pairwise and single-sample populations differ, and training exposure varies across model groups. These observations describe rank instability; they do not isolate causal effects of format or model size.

The third finding is about the difference between error and information. A regressor that predicts near the centre of the target distribution can achieve a lower MAE than a model that attempts to track the target, because absolute error is forgiving of central predictions when the target is bounded. Within the common archived fibrosis set, Qwen3-1.7B v5 has conditional MAE 1.142 (162/162 parsed) with ρ = 0.105, while Llama 3.3 70B has conditional MAE 1.517 (150/162 parsed) with ρ = 0.017; on the 150 jointly parsed items the MAEs are 1.127 and 1.517. This is a descriptive contrast, not a model-class superiority test. Low MAE alone does not show that a model distinguishes fibrosis severity between samples. Predictions varied less than the observed stages and had weak rank correlations with them; the reason for this pattern remains uncertain.

For cfRNA, second cfRNA cohort accuracy is 0.620 [0.553, 0.687] under the representation calculated from all samples, including held-out samples, 0.609 [0.536, 0.676] with training-only gene-wise preprocessing and 0.598 [0.531, 0.665] with per-sample standardisation. The first two lower bounds exceed the observed majority of 0.531; the third does not. The 1.1-percentage-point difference between the latter point estimates is not a demonstrated between-pipeline difference. The reference cohort analyses do not demonstrate an accuracy advantage over their majority baseline of 0.714 under these procedures. The small cohorts and wide intervals limit interpretation.

Reader disagreement depends on the outcome and model. The two readers assign the same stage to 56 of 108 jointly assessed samples; unweighted κ is 0.400 and quadratic-weighted κ is 0.809. Changing the reference reader changes binary accuracy by at most 4.6 percentage points across the evaluated archives and five-class accuracy by 13 percentage points for one classical model. These differences neither establish a universal limit on model performance nor identify either reader as incorrect.

Finally, the operational record matters. One system produced no valid cfRNA completion because its requests failed; this does not describe its evaluations in other families. Several others produced parseable answers for a small minority of items. Reporting those zeros as capability would have been a factual error about the models; excluding them silently would have been a factual error about the evaluation. The benchmark’s value depends on keeping those categories separate.

## 5. Limitations

1. **Small cohorts and few positives.** The cfRNA reference cohort has 112 samples and 32 positives; the second cfRNA cohort has 179 and 84. Confidence intervals are wide, and a 112-sample cohort cannot reliably distinguish a modestly better-than-majority classifier from the majority classifier itself. The tissue cohorts are similarly bounded at 168 profiles.
2. **Tissue train–test overlap.** The frozen MB-20xx tissue test set and the MB-12xx staged-liver SFT pool both derive from GSE135251, and the v6/v7 checkpoints were trained on the test samples (Methods). They are therefore excluded from the primary tissue rankings and generalization claims and reported only as diagnostics. The builder script for MB-12xx is not preserved in either repository, and the original EPoS/Kamzolas source files are absent locally; accession identity is supported by GEO identifiers and expression agreement, but the complete sequence of source files and processing steps remains a reproducibility gap.
3. **Evaluated populations and dependence.** Response archives do not contain prompt text, so prompt-byte identity cannot be verified from the archives; the validated join is task, sample identifier and reference label. Some models were queried on different item sets, especially for pairwise tasks. The primary single-sample scores use common item intersections, and pairwise results are reported within exact recorded evaluation sets. A biological-sample cluster bootstrap is used for pairwise intervals, but this does not capture every source of dependence; broader patient/donor-clustered inference and a preregistered recorded evaluation set remain necessary before strengthening any inferential claim.
4. **Representation dependence.** Primary frozen-corpus classical estimates are transductive. Training-only gene-wise preprocessing gives second cfRNA cohort accuracy 0.609 [0.536, 0.676] and reference cohort accuracy 0.714 [0.634, 0.795]. The second cfRNA cohort confidence-interval criterion is not robust to all tested representations. None of these retrospective estimates establishes prospective clinical performance.
5. **Incomplete coverage and truncation.** Parsed-answer coverage varies from 0% to 100% across models and tasks. The cfRNA evaluation script used a 900-token output budget rather than the planned 1,500, and compact evaluation used 24 tokens. Truncation is a property of the evaluation software, not of the model, and it depresses measured accuracy for verbose reasoners.
6. **Contamination cannot be excluded retrospectively.** The datasets are built from public cohorts that may appear in pretraining data. No canary probe was planted, and a probe planted now could not retrospectively certify the archives. Memorisation testing is future work.
7. **Registration is retrospective.** Corpora were frozen and checksummed before evaluation, and the correction log is versioned, but the freeze is a snapshot rather than a prospective registration. It does not eliminate analytic flexibility, and it does not establish that the analysis plan preceded data access.
8. **Single-family fine-tuning.** The compact arm is a Qwen3 family with one quantised 8B run. Comparisons between compact and API-accessed model groups confound model class with training exposure. No claim about scaling behaviour in general follows from this model set.
9. **No clinical comparator.** The tested inputs are molecular measurements only. Routine clinical covariates and validated scores such as FIB-4, APRI and NFS were deliberately excluded. The benchmark therefore cannot show superiority over clinical practice, and nothing here supports screening, diagnosis, treatment monitoring or triage.
10. **Clinical-consensus labels in the second cfRNA cohort.** Reference labels in this cohort are based on clinical consensus; biopsy assessments are available for only a 42-patient subset. The labels are not based on adjudicated readings by two biopsy readers for every sample. Clinical-consensus labels are not equivalent to histological assessments.
11. **Visit selection is order-dependent.** The second cfRNA cohort picks the first Baseline row per patient with a first-row fallback. The number of affected patients was not quantified.
12. **Batch checks are incomplete.** A non-significant association between fibrosis and sequencing run (χ² = 17.9, p = 0.332) is not evidence that confounding is absent. The planned hospital-level check was not implemented.
13. **Provenance gaps.** The evaluation script archived the requested model string but not a provider-returned identifier, usage counters or finish reasons, so a frozen provider snapshot cannot be demonstrated from the archive alone.

## 6. Conclusions

MASLD-Bench evaluates instruction-following language models on molecular inputs using explicit task definitions and evaluated populations. On the aligned tissue binary task, none of the 17 eligible models exceeds the majority accuracy; checkpoints with documented test-set exposure are excluded. Conditional regression error must be interpreted alongside output coverage and rank correlation. Rank differences across answer formats are descriptive rather than causal estimates of format or model-size effects.

In cfRNA, the second cfRNA cohort logistic result meets a specified confidence-interval criterion under training-only gene-wise preprocessing but not under every representation. The reference cohort analyses do not demonstrate an accuracy advantage over the reference cohort majority baseline of 0.714 under the evaluated procedures. The external five-gene result is exploratory: its accuracy is descriptively above the second cfRNA cohort majority of 0.531, but its paired comparison is not significant. These findings do not establish a validated molecular signature, an absence of biological information, or clinical readiness. This does not mean language models are incapable of assisting with MASLD. Rather, it means that current zero-shot prompting on raw molecular panels is insufficient. Future AI systems will likely require specialized architectures that natively understand genomic coordinates and biological pathways, rather than treating genes as strings of text

## 7. Declarations and availability

### Author contributions

Subhadip Banerjee: Conceptualization, Methodology, Software, Investigation, Data curation, Formal analysis, Visualization, Writing – original draft. Wim Vanden Berghe: Conceptualization, Formal analysis, Writing – review & editing, Supervision. Rawiwan Charoensub: Conceptualization, Resources, Supervision, Project administration, Writing – review & editing.

### Ethics and consent

This study reanalysed existing datasets and collected no new biological material.

### Data and code availability

The benchmark package is archived on Zenodo under concept DOI 10.5281/zenodo.23836159; the current version is 10.5281/zenodo.23855439 (notebooks, site, run log; corrected 2026-11-01; access restricted until publication). The plasma cfRNA source cohort is 10.5281/zenodo.19047046 (Poma-Soto et al.). Gated cfRNA data are available through EGA under accession EGAD50000000775 / study EGAS50000000545 under the Ghent DAC Data Usage Agreement; per-sample values and prompt texts are not redistributed. Fine-tuned checkpoints are deposited at Hugging Face under the author’s account: `Subhadip-igtm/masld-bench-qwen3-8b-sft-v7` and `Subhadip-igtm/masld-bench-qwen3-8b-sft-v7-adapter`; `Subhadip-igtm/masld-bench-qwen3-1.7b-sft-v6` and `Subhadip-igtm/masld-bench-qwen3-1.7b-sft-v6-adapter` (https://huggingface.co/Subhadip-igtm); each model card carries the Zenodo, OSF and EGA identifiers and the Apache-2.0 license. OSF registration (retrospective deposit of the frozen analysis plans and run log): https://osf.io/4vhbs/. Current and historical leaderboard standings for all tasks and models are available in the benchmark package; a public web leaderboard is scaffolded but not yet deployed.

## Declaration of generative AI and AI-assisted technologies in the manuscript preparation process

The authors report using GLM 5.3, Deepseek 3.0 Pro, GPT-6 Astra and Kimi 3 pro. Codex was also used to repair and edit manuscript text, prepare document-building code and check the resulting files. Additionally, generative AI tools were used to assist in the conceptualization and drafting of the workflow schematic in Figure 1. The tool used for Figure 1 was GPT-6 Astra (OpenAI).

## Acknowledgements

The Ghent cohort was generated by the Ghent University research groups responsible for the plasma cell-free RNA biobank and cohort, who act as the Ghent DAC for the EGA deposition (EGAD50000000775 / EGAS50000000545); we thank them for making the cohort available under the Data Usage Agreement and for the annotation that made the construction of the biopsy-referenced dataset possible. S.B. acknowledges the Research Foundation – Flanders (FWO), which allocated a grant for a scientific stay in Flanders (file ID V504225N, hosted at the University of Antwerp) during which part of this work was done. R.C. acknowledges the Ursimone Wietlisbach Foundation for grant support.

## Supplementary material for MASLD-Bench

**Supplementary Table S1.**
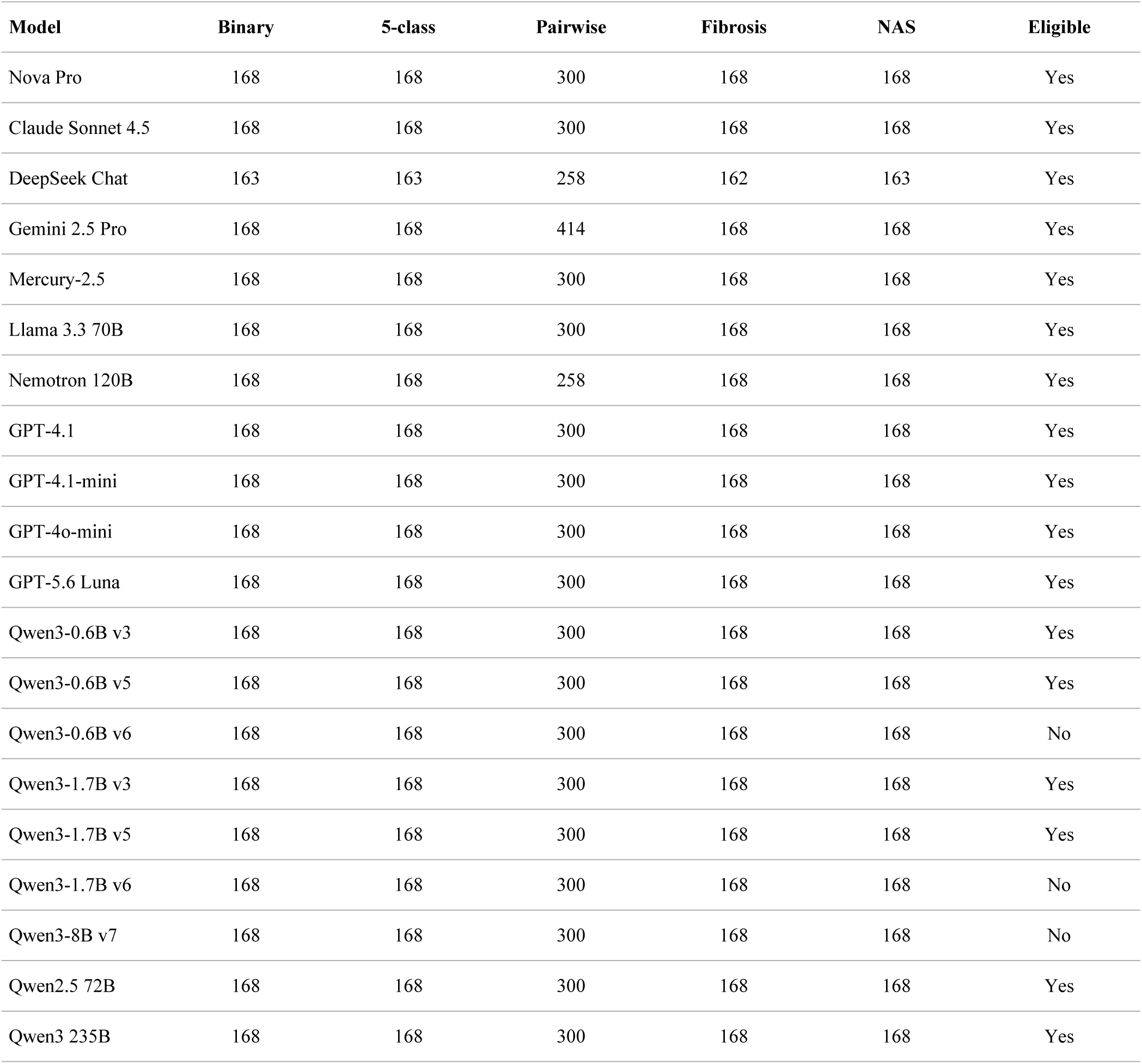
Archive coverage before alignment. Counts below are archived in-corpus items, not the final common-item denominators. Columns identify binary, five-class, pairwise, fibrosis-regression and NAS tasks. Three exposed checkpoints are excluded from primary tissue comparisons. The full audit TSV includes duplicate keys, superseded rows and gold mismatches.

**Supplementary Table S2.**
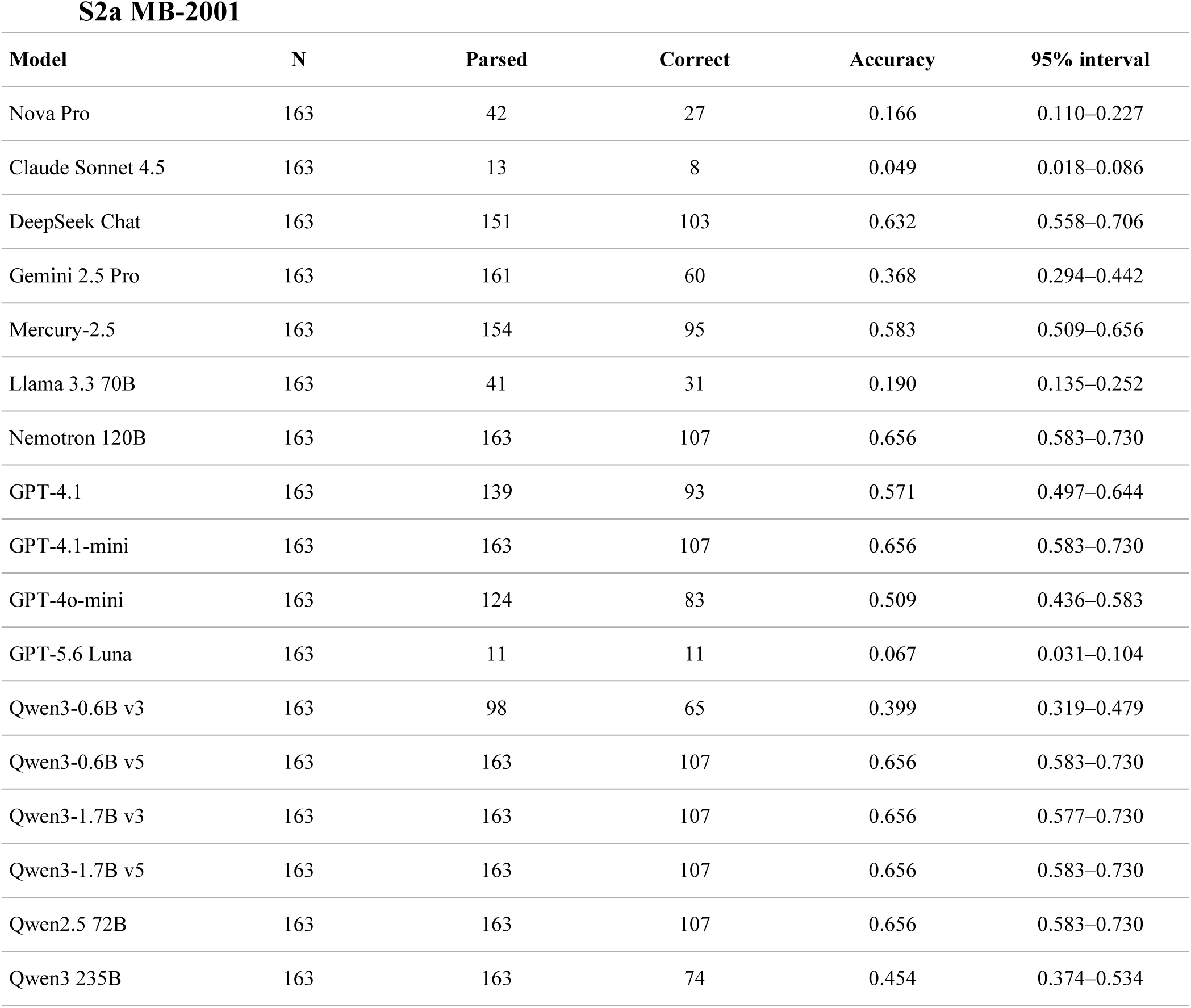
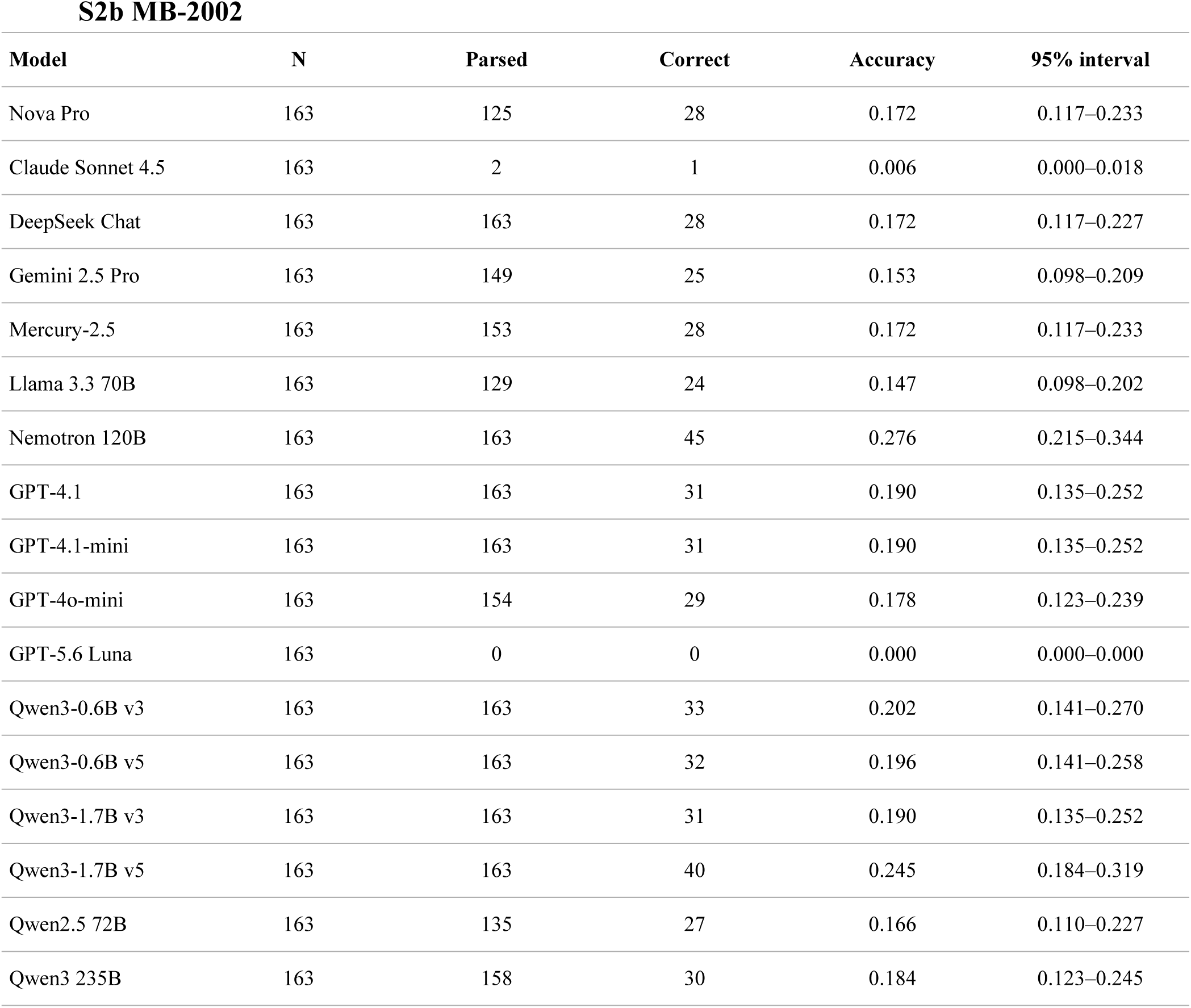
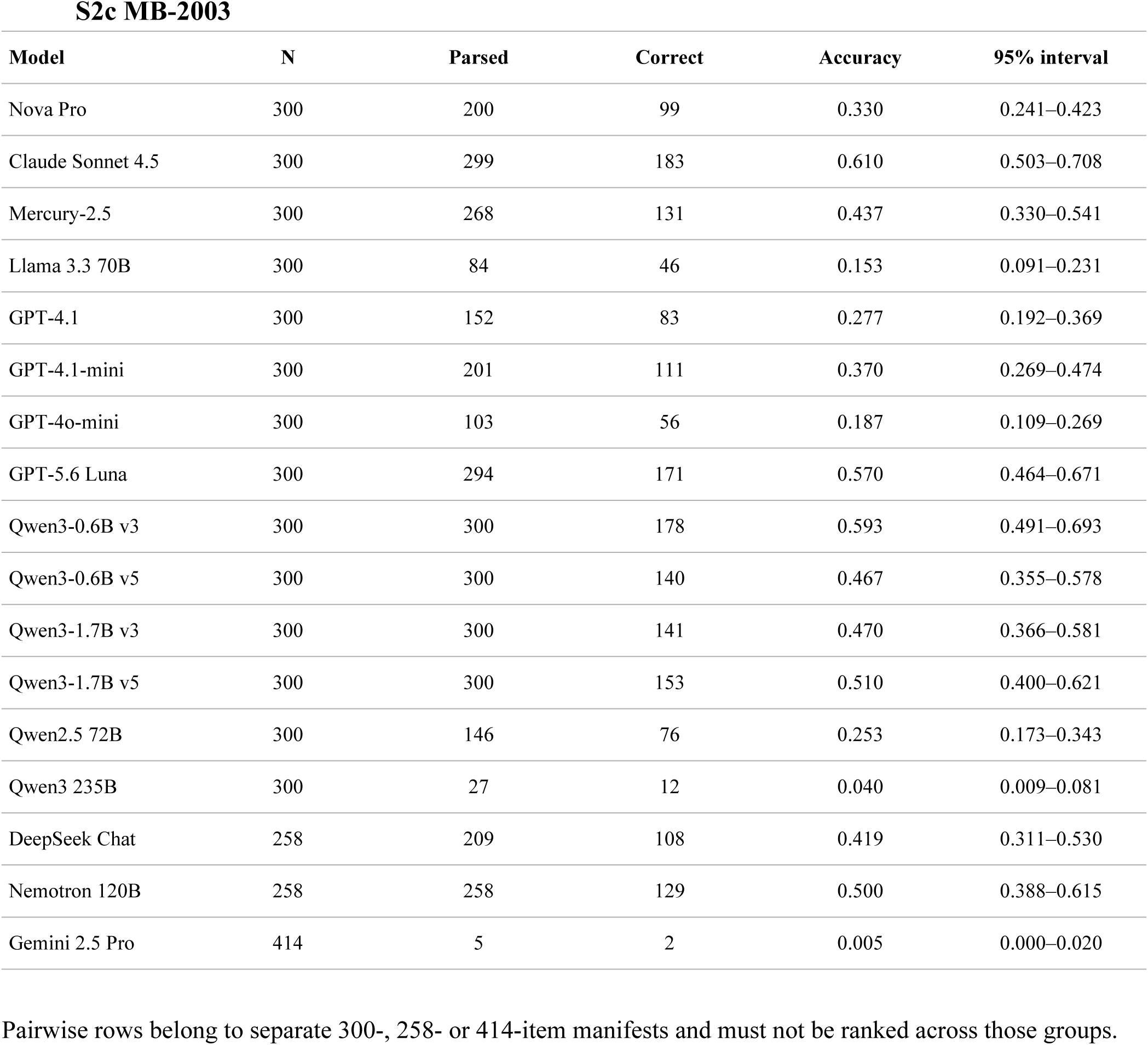
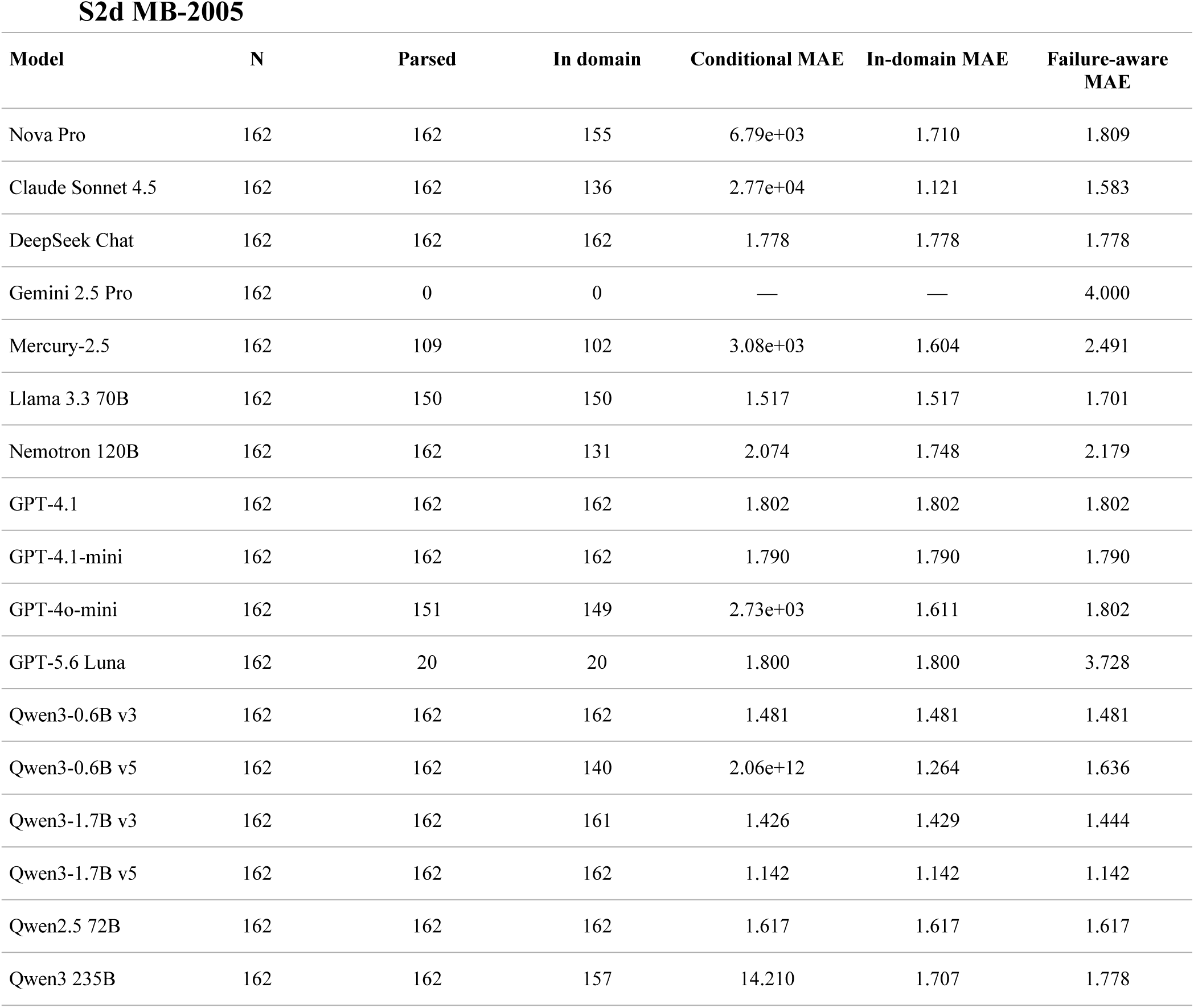
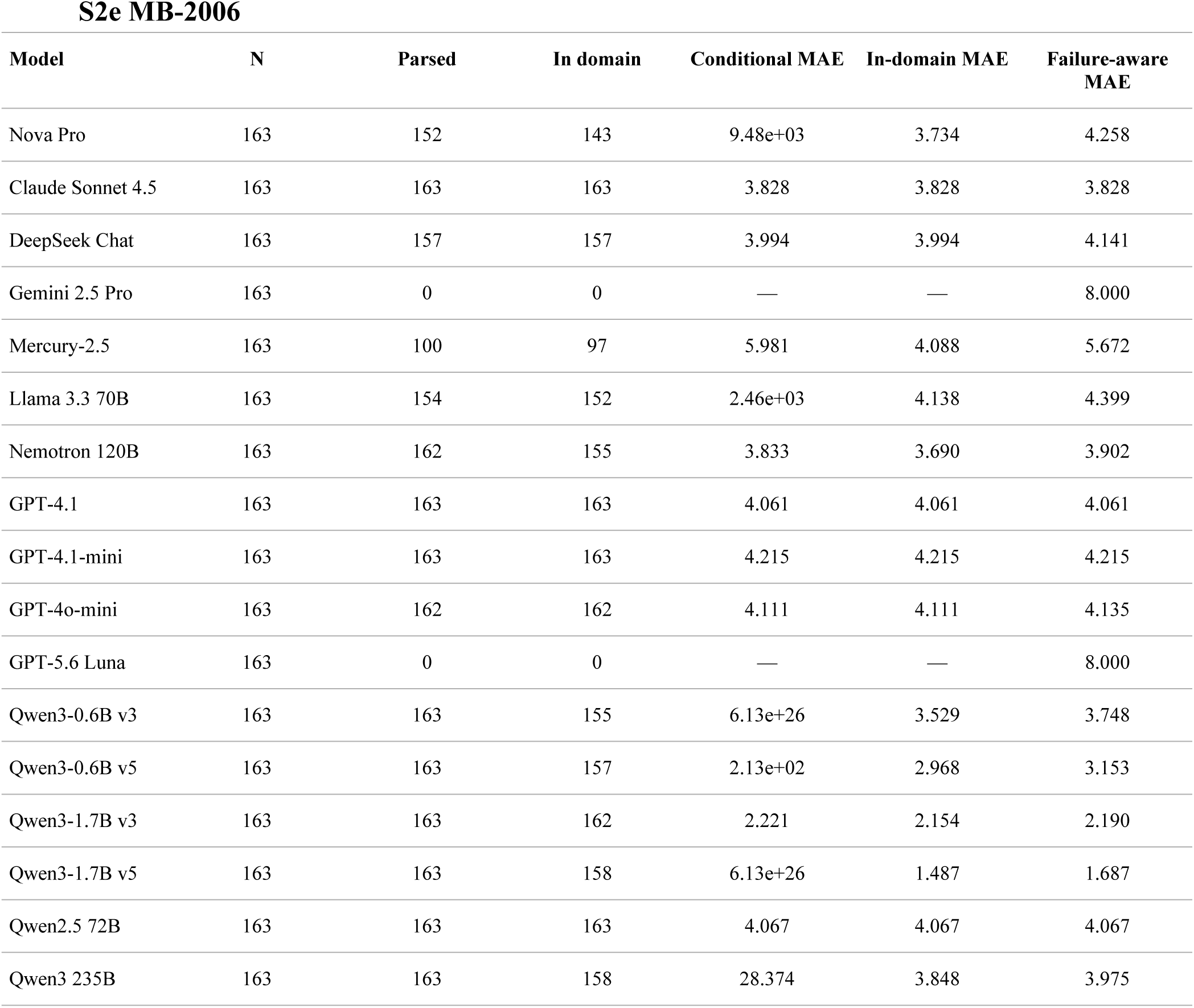
Coverage and regression sensitivity for the eligible tissue roster. Conditional MAE uses parsed numeric outputs, including out-of-domain values. In-domain MAE uses parsed outputs within fibrosis 0–4 or NAS 0–8. Failure-aware MAE averages over all evaluated items, retaining in-domain absolute errors and assigning a penalty of 4 (fibrosis) or 8 (NAS) otherwise. This policy is a declared sensitivity, not imputation. Classification counts invalid answers as errors.

**Figure S1.**
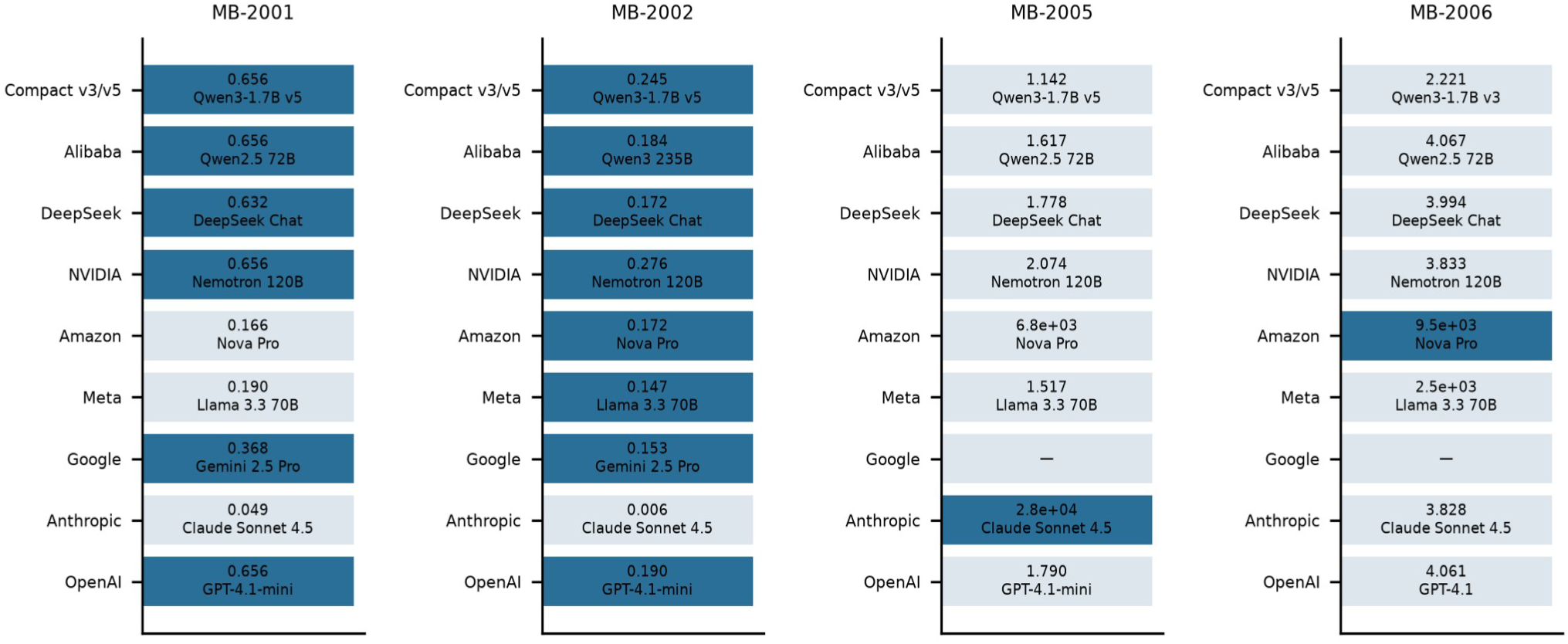
Best eligible model within each developer/group on the four aligned single-sample tissue tasks: binary advanced fibrosis (MB-2001), five-class fibrosis stage (MB-2002), numeric fibrosis stage (MB-2005), and numeric NAFLD Activity Score (MB-2006). Model selection is descriptive and performed on these evaluation sets. Pairwise results are omitted because manifests differ. Values are accuracy for classification and conditional MAE for regression; extreme values indicate numeric-format failures.

**Figure S2.**
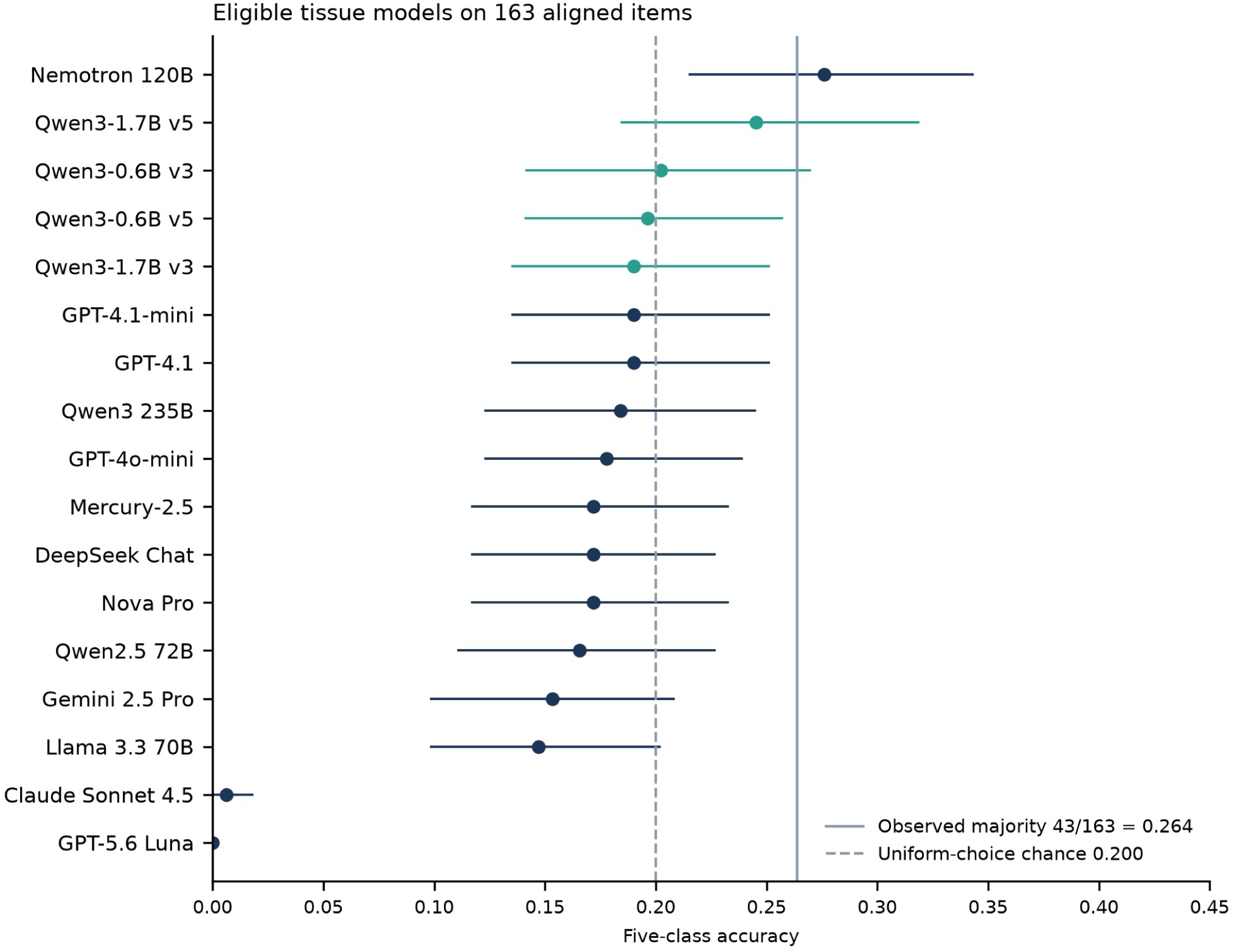
Five-class accuracy for 17 eligible tissue models on 163 aligned items. The solid line is observed majority 43/163 = 0.264 and the dashed line uniform-choice chance 0.200. Historical blood-proxy and plasma panels are excluded because they were not re-derived on the authoritative path.

**Figure S3.**
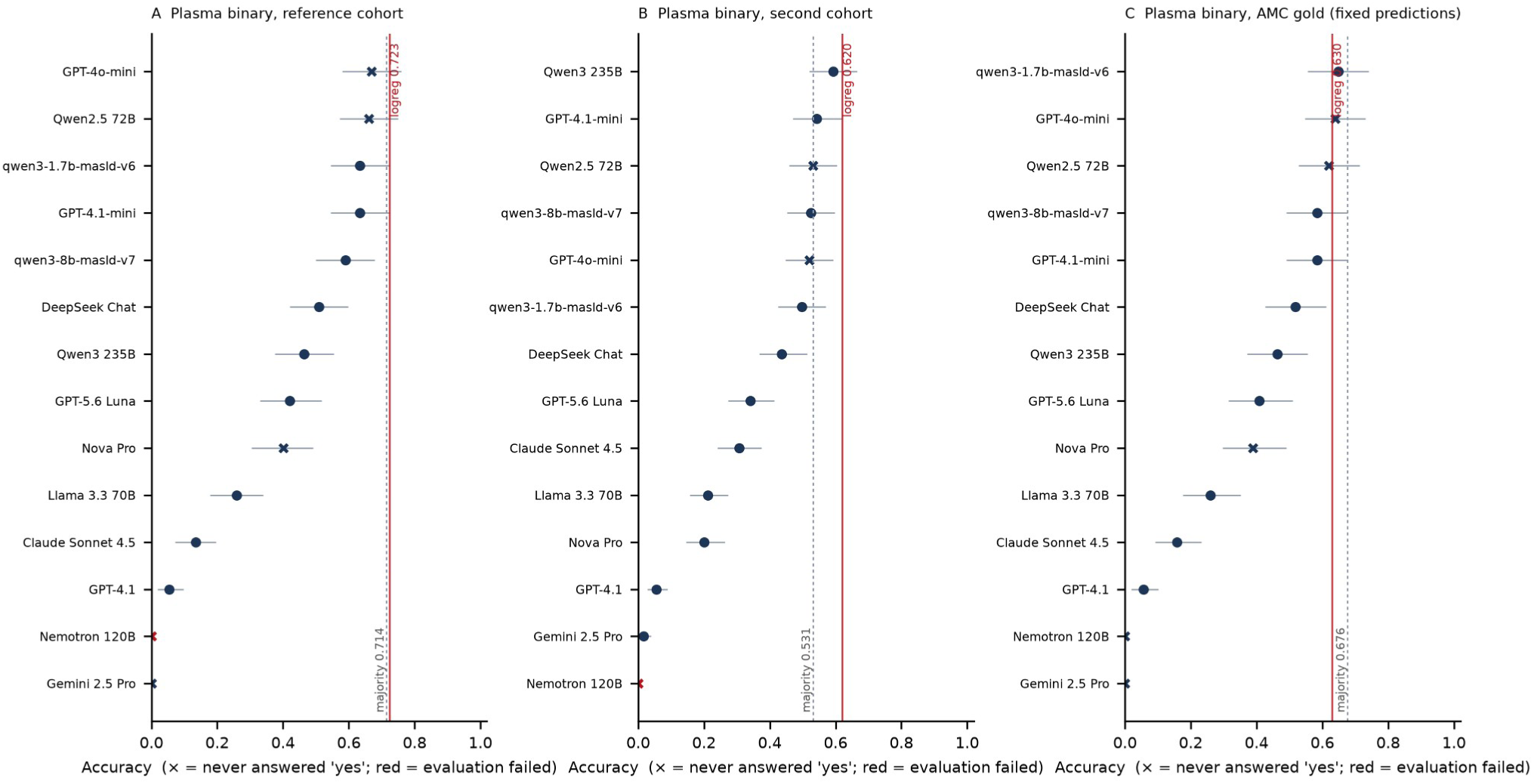
Plasma binary results on reference-cohort, second-cohort and AMC-reader labels. Dashed majority lines are 0.714, 0.531 and 0.676. Red classical references are 0.723, 0.620 and 0.630. AMC uses fixed reference-cohort model predictions on 108 samples; it is a reader sensitivity, not an independent external cohort.

**Figure S4.**
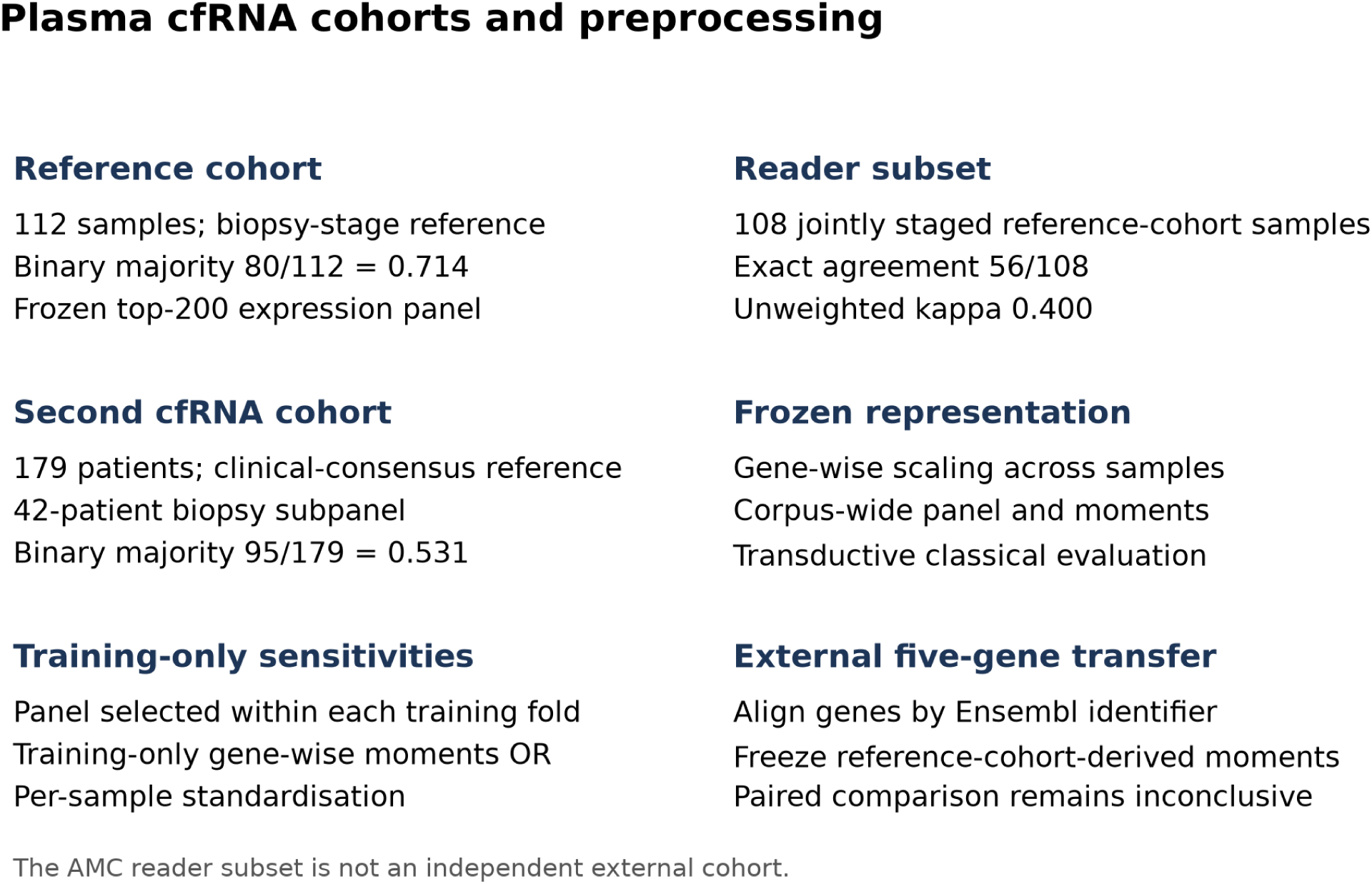
Cohorts and preprocessing contracts for cfRNA. The frozen representation uses gene-wise scaling across samples. Training-only sensitivities and external transfer are distinct. Reader agreement is based on 108 jointly staged samples.

**Figure S5.**
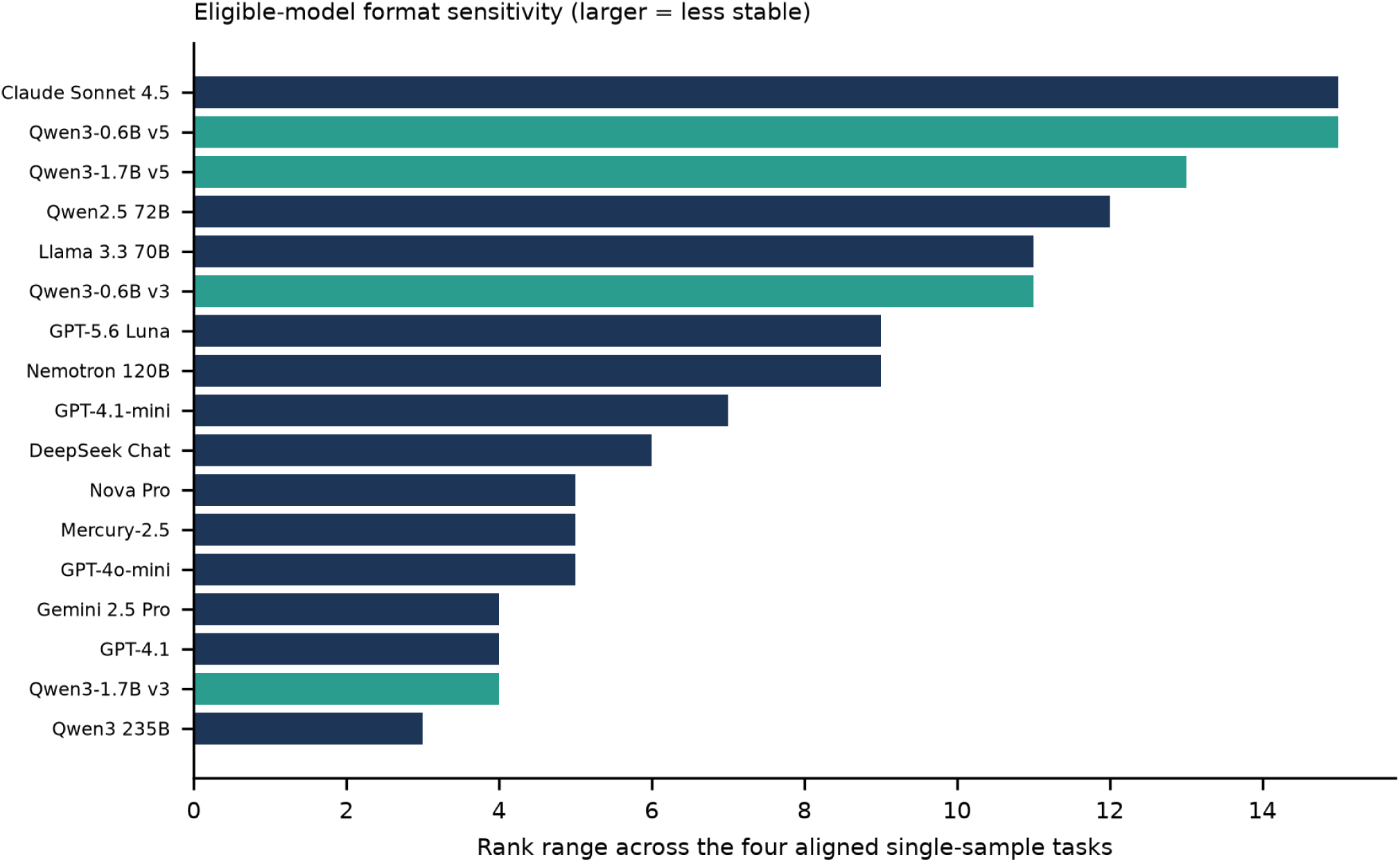
Within-model rank range over the four aligned single-sample tissue tasks. Pairwise ranks are excluded. Larger ranges indicate greater descriptive variation across task formats; they do not isolate format effects.

**Figure S6.**
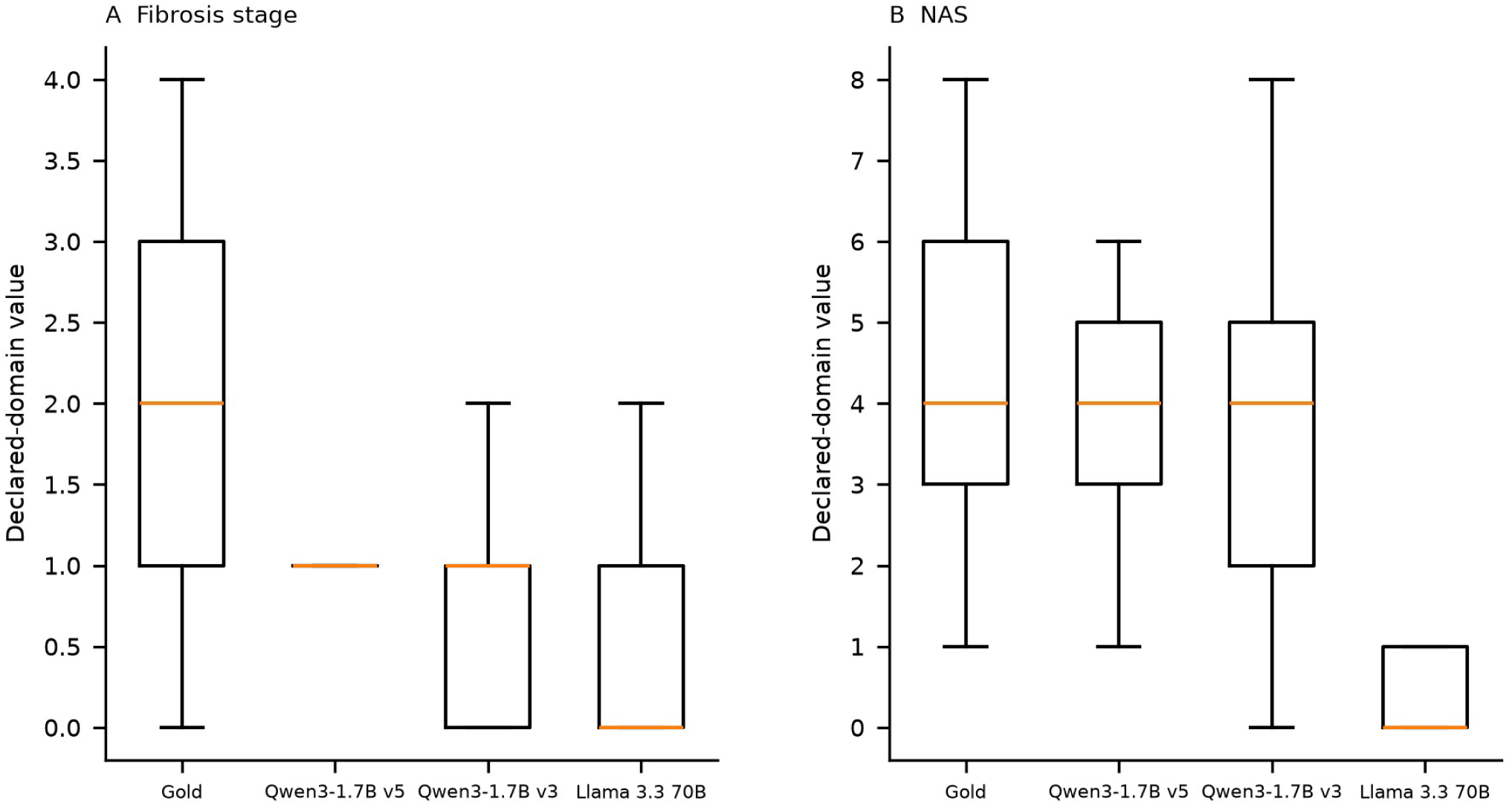
Gold and predicted distributions for selected eligible tissue models. Predictions are restricted to the declared domains (fibrosis 0–4; NAS 0–8). Boxes show interquartile ranges and median; whiskers follow the plotting library default and outliers are hidden. These parsing- and domain-selected distributions do not replace whole-set failure-aware results.

**Figure S7.**
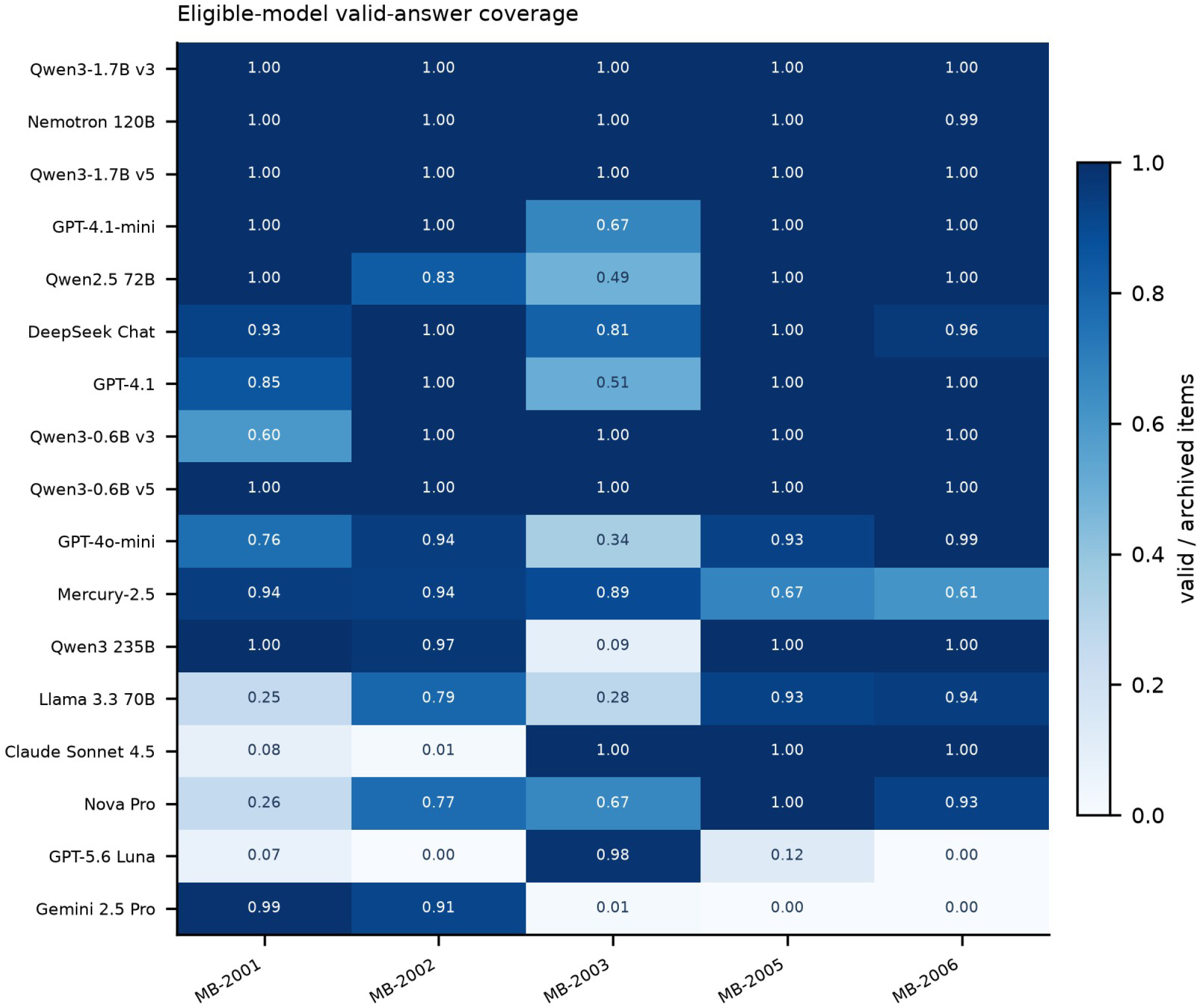
Parsed-answer coverage on aligned single-sample tasks and the respective pairwise manifests. Values are valid/archived items. Coverage is distinct from correctness and, for regression, from domain validity.

**Figure S8.**
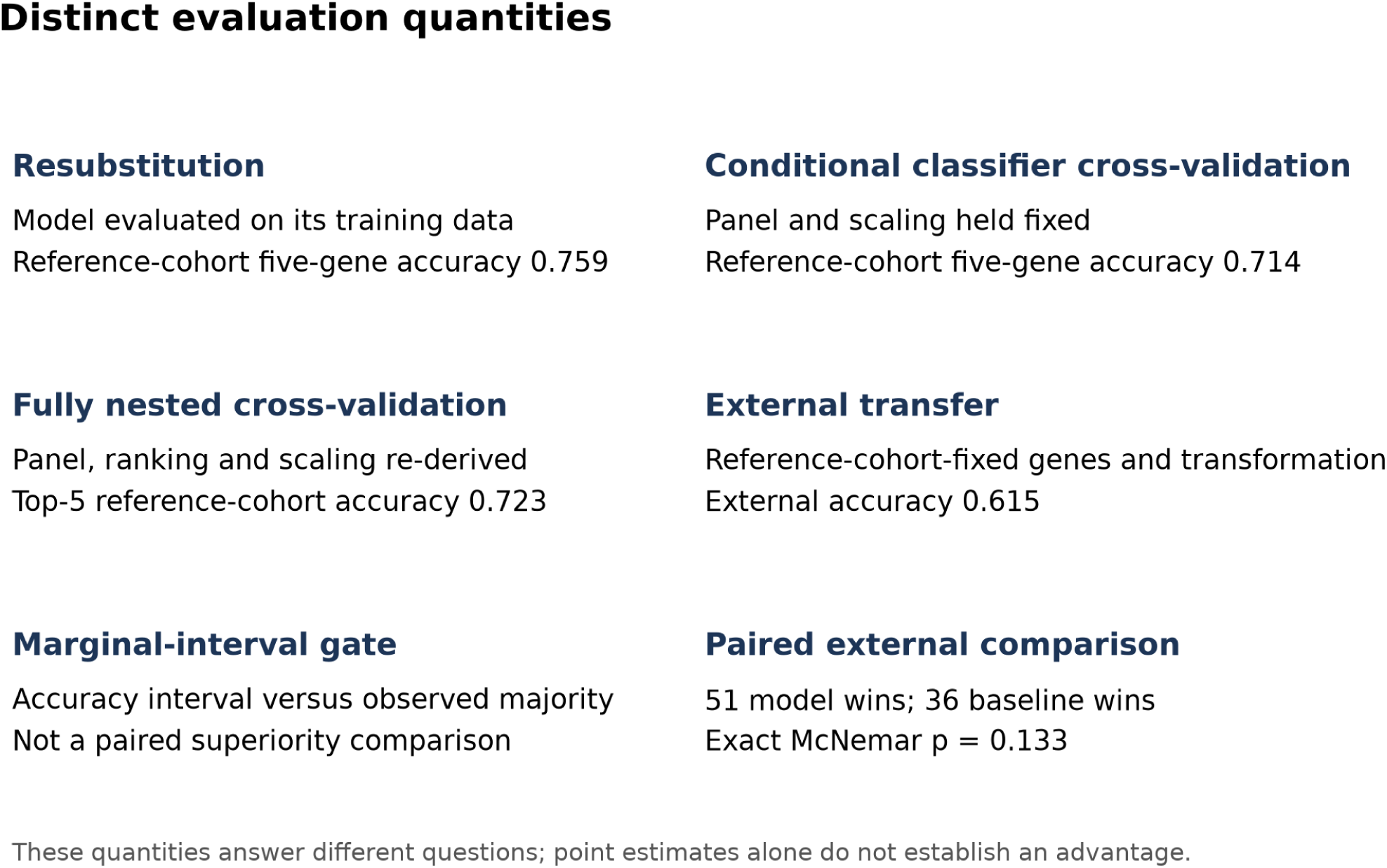
Interpretation of distinct cfRNA validation quantities. Resubstitution, fixed-panel classifier cross-validation, fully nested cross-validation and external transfer evaluate different quantities. The external paired comparison does not establish superiority.

**Figure S9.**
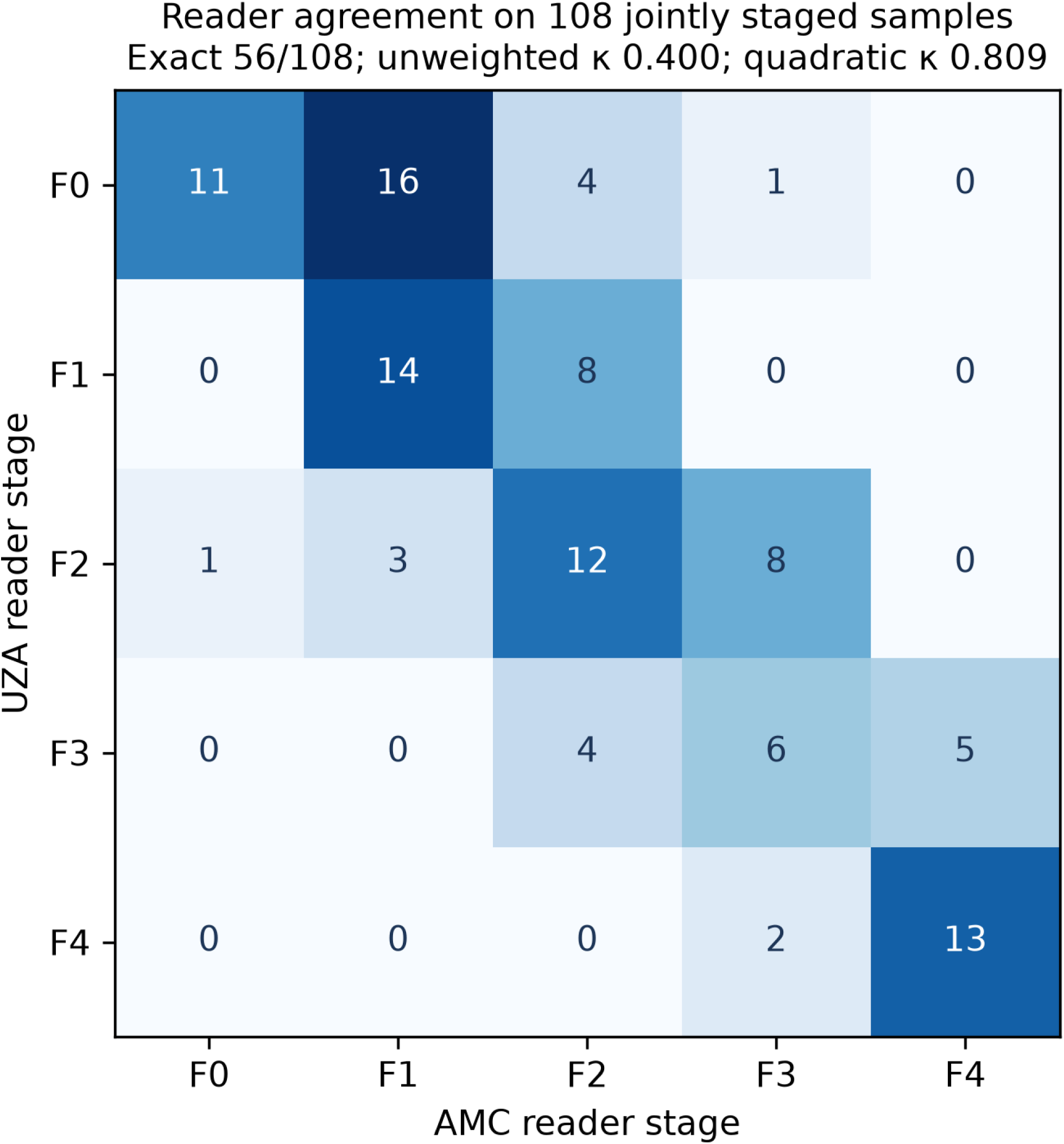
Confusion matrix between UZA and AMC biopsy readings for 108 jointly staged samples. Exact agreement is 56/108, unweighted κ = 0.400 and quadratic-weighted κ = 0.809. The aggregate matrix is reproduced from the corrected reader-agreement figure; no patient-level identifiers are included.

## References

[1] Younossi ZM, Koenig AB, Abdelatif D, Fazel Y, Henry L, Wymer M. Global epidemiology of nonalcoholic fatty liver disease—Meta analytic assessment of prevalence, incidence, and outcomes. Hepatology. 2016;64(1):73–84. doi:10.1002/hep.28431.

[2] Chew NW, Ng CH, Tan DJH, Kong G, Lin C, Chin YH, et al. The global burden of metabolic disease: Data from 2000 to 2019. Cell Metab. 2023;35(3):414–428.e3. doi:10.1016/j.cmet.2023.02.003.

3. European Association for the Study of the Liver (EASL), European Association for the Study of Diabetes (EASD), European Association for the Study of Obesity (EASO). EASL-EASD-EASO Clinical Practice Guidelines on the Management of Metabolic Dysfunction-Associated Steatotic Liver Disease (MASLD). Obes Facts. 2024;17(4):374-444. doi:10.1159/000539371.

[4] Kleiner DE, Brunt EM, Van Natta M, Behling C, Contos MJ, Cummings OW, et al. Design and validation of a histological scoring system for nonalcoholic fatty liver disease. Hepatology. 2005;41(6):1313–1321. doi:10.1002/hep.20701.

[5] Abdelhameed F, Kite C, Lagojda L, Dallaway A, Chatha KK, Chaggar SS, et al. Non-invasive Scores and Serum Biomarkers for Fatty Liver in the Era of Metabolic Dysfunction-associated Steatotic Liver Disease (MASLD): A Comprehensive Review From NAFLD to MAFLD and MASLD. Curr Obes Rep. 2024;13(3):510–531. doi:10.1007/s13679-024-00574-z.

[6] Zhavoronkov A, Naumov V, Sidorenko D, Aliper A, Aladinskiy V, Hasani R, et al. An open benchmark and language models for AI in aging biology. Cell. 2026;189(19):5980–5994.e8. doi:10.1016/j.cell.2026.08.026.

[7] Zhavoronkov A, Sidorenko D, Naumov V, Pushkov S, Zagirova D, Aladinskiy V, et al. LongevityBench: Are SotA LLMs ready for aging research?. bioRxiv preprint. 2026. doi:10.64898/2026.01.12.698650.

[8] Chen SF, Alyakin A, Seas A, Yang E, Choi JJ, Lee JV, et al. LLM-assisted systematic review of large language models in clinical medicine. Nat Med. 2026;32(3):1152–1159. doi:10.1038/s41591-026-04229-5.

[9] Njei B, Al Ajlouni YA, Ajayi A, Shahin F, Ta’ani OA, Boateng S, et al. Large Language Models for Diagnosis and Prognosis of Chronic Liver Diseases: A Systematic Review. Health Sci Rep. 2026;9(5):e72476. doi:10.1002/hsr2.72476.

[10] Njei B, Njei N, Boateng S, Al Ta’ani O, Al-Ajlouni Y. Using Large Language Models to Predict Advanced Liver Fibrosis in Metabolic Dysfunction-Associated Steatotic Liver Disease (MASLD): A Proof-of-Concept Analysis. Cureus. 2026. doi:10.7759/cureus.102777.

[11] Wu W, Guo Y, Li Q, Jia C. Exploring the potential of large language models in identifying metabolic dysfunction associated steatotic liver disease: A comparative study of non invasive tests and artificial intelligence generated responses. Liver Int. 2025;45(4):e16112. doi:10.1111/liv.16112.

[12] Pugliese N, Giuffrè M, Schattenberg JM. Large Language Models in MASLD: The New Era of Generative Artificial Intelligence Augmented Clinical Practice. Liver Int. 2025;45(4):e16162. doi:10.1111/liv.16162.

[13] Paverd H, Gao Z, Mahani G, Fabre MA, Burge S, Hoare M, et al. LLM-based reconstruction of longitudinal clinical trajectories in chronic liver disease. npj Precis Oncol. 2026. doi:10.1038/s41698-026-01650-4.

[14] Wang Y, Fan Y, Wang X, Yu T, Zong Y, Liu X, et al. SCMBench: benchmarking domain-specific and foundation models for single-cell multi-omics data integration. Nat Commun. 2026;17(1):5967. doi:10.1038/s41467-026-72570-x.

[15] Bendidi I, Whitfield S, Kenyon-Dean K, Yedder HB, Mesbahi YE, Noutahi E, et al. Benchmarking Transcriptomics Foundation Models for Perturbation Analysis : one PCA still rules them all. arXiv preprint. 2024. doi:10.48550/arXiv.2410.13956.

[16] Shaked G, Nagisetty S. OmicsBench: a long-horizon multi-turn benchmark of AI agents on research-study-scale bioinformatics. arXiv preprint. 2026. doi:10.48550/arXiv.2610.21074.

[17] Luo E, Jia J, Xiong Y, Li X, Guo X, Yu B, et al. Benchmarking AI scientists for omics data–driven biological discovery. Bioinformatics. 2026;42(Supplement_1):btag227. doi:10.1093/bioinformatics/btag227.

[18] Dietterich TG. Approximate Statistical Tests for Comparing Supervised Classification Learning Algorithms. Neural Comput. 1998;10(7):1895–1923. doi:10.1162/089976698300017197.

19. Wu S, Nair Y, Candès EJ. Efficient Evaluation of LLM Performance with Statistical Guarantees. arXiv preprint. 2026. doi:10.48550/arXiv.2601.20251.

[20] Ghosh S, Zhao X, Alim M, Brudno M, Bhat M. Artificial intelligence applied to ‘omics data in liver disease: towards a personalised approach for diagnosis, prognosis and treatment. Gut. 2025;74(2):295–311. doi:10.1136/gutjnl-2023-331740.

[21] Govaere O, Cockell S, Tiniakos D, Queen R, Younes R, Vacca M, et al. Transcriptomic profiling across the nonalcoholic fatty liver disease spectrum reveals gene signatures for steatohepatitis and fibrosis. Sci Transl Med. 2020;12(572):eaba4448. doi:10.1126/scitranslmed.aba4448.

[22] Kamzolas I, Koutsandreas T, Barker CG, Vathrakokoili Pournara A, Weston H, Fujiwara N, et al. A data-driven framework reconstructs the molecular continuum of human MASLD progression. Nat Metab. 2026;8(7):1545–1562. doi:10.1038/s42255-026-01543-7.

