## Supplementary Table and Figure for "MASLD-Bench evaluating language models on liver omics and fibrosis endpoints"

### Supplementary material for MASLD-Bench

#### Supplementary Table S1

Archive coverage before alignment. Counts below are archived in-corpus items, not the final common-item denominators. Columns identify binary, five-class, pairwise, fibrosis-regression and NAS tasks. Three exposed checkpoints are excluded from primary tissue comparisons. The full audit TSV includes duplicate keys, superseded rows and gold mismatches.

| Model | Binary | 5-class | Pairwise | Fibrosis | NAS | Eligible |
|---|---|---|---|---|---|---|
| Nova Pro | 168 | 168 | 300 | 168 | 168 | Yes |
| Claude Sonnet 4.5 | 168 | 168 | 300 | 168 | 168 | Yes |
| DeepSeek Chat | 163 | 163 | 258 | 162 | 163 | Yes |
| Gemini 2.5 Pro | 168 | 168 | 414 | 168 | 168 | Yes |
| Mercury-2.5 | 168 | 168 | 300 | 168 | 168 | Yes |
| Llama 3.3 70B | 168 | 168 | 300 | 168 | 168 | Yes |
| Nemotron 120B | 168 | 168 | 258 | 168 | 168 | Yes |
| GPT-4.1 | 168 | 168 | 300 | 168 | 168 | Yes |
| GPT-4.1-mini | 168 | 168 | 300 | 168 | 168 | Yes |
| GPT-4o-mini | 168 | 168 | 300 | 168 | 168 | Yes |
| GPT-5.6 Luna | 168 | 168 | 300 | 168 | 168 | Yes |
| Qwen3-0.6B v3 | 168 | 168 | 300 | 168 | 168 | Yes |
| Qwen3-0.6B v5 | 168 | 168 | 300 | 168 | 168 | Yes |
| Qwen3-0.6B v6 | 168 | 168 | 300 | 168 | 168 | No |
| Qwen3-1.7B v3 | 168 | 168 | 300 | 168 | 168 | Yes |
| Qwen3-1.7B v5 | 168 | 168 | 300 | 168 | 168 | Yes |
| Qwen3-1.7B v6 | 168 | 168 | 300 | 168 | 168 | No |
| Qwen3-8B v7 | 168 | 168 | 300 | 168 | 168 | No |
| Qwen2.5 72B | 168 | 168 | 300 | 168 | 168 | Yes |
| Qwen3 235B | 168 | 168 | 300 | 168 | 168 | Yes |

#### Supplementary Table S2

Coverage and regression sensitivity for the eligible tissue roster. Conditional MAE uses parsed numeric outputs, including out-of-domain values. In-domain MAE uses parsed outputs within fibrosis 0–4 or NAS 0–8. Failure-aware MAE averages over all evaluated items, retaining in-domain absolute errors and assigning a penalty of 4 (fibrosis) or 8 (NAS) otherwise. This policy is a declared sensitivity, not imputation. Classification counts invalid answers as errors.

#### Table S2a MB-2001

| Model | N | Parsed | Correct | Accuracy | 95% interval |
|---|---|---|---|---|---|
| Nova Pro | 163 | 42 | 27 | 0.166 | 0.110–0.227 |
| Claude Sonnet 4.5 | 163 | 13 | 8 | 0.049 | 0.018–0.086 |
| DeepSeek Chat | 163 | 151 | 103 | 0.632 | 0.558–0.706 |
| Gemini 2.5 Pro | 163 | 161 | 60 | 0.368 | 0.294–0.442 |
| Mercury-2.5 | 163 | 154 | 95 | 0.583 | 0.509–0.656 |
| Llama 3.3 70B | 163 | 41 | 31 | 0.190 | 0.135–0.252 |
| Nemotron 120B | 163 | 163 | 107 | 0.656 | 0.583–0.730 |
| GPT-4.1 | 163 | 139 | 93 | 0.571 | 0.497–0.644 |
| GPT-4.1-mini | 163 | 163 | 107 | 0.656 | 0.583–0.730 |
| GPT-4o-mini | 163 | 124 | 83 | 0.509 | 0.436–0.583 |
| GPT-5.6 Luna | 163 | 11 | 11 | 0.067 | 0.031–0.104 |
| Qwen3-0.6B v3 | 163 | 98 | 65 | 0.399 | 0.319–0.479 |
| Qwen3-0.6B v5 | 163 | 163 | 107 | 0.656 | 0.583–0.730 |
| Qwen3-1.7B v3 | 163 | 163 | 107 | 0.656 | 0.577–0.730 |
| Qwen3-1.7B v5 | 163 | 163 | 107 | 0.656 | 0.583–0.730 |
| Qwen2.5 72B | 163 | 163 | 107 | 0.656 | 0.583–0.730 |
| Qwen3 235B | 163 | 163 | 74 | 0.454 | 0.374–0.534 |

#### Table S2b MB-2002

| Model | N | Parsed | Correct | Accuracy | 95% interval |
|---|---|---|---|---|---|
| Nova Pro | 163 | 125 | 28 | 0.172 | 0.117–0.233 |
| Claude Sonnet 4.5 | 163 | 2 | 1 | 0.006 | 0.000–0.018 |
| DeepSeek Chat | 163 | 163 | 28 | 0.172 | 0.117–0.227 |
| Gemini 2.5 Pro | 163 | 149 | 25 | 0.153 | 0.098–0.209 |
| Mercury-2.5 | 163 | 153 | 28 | 0.172 | 0.117–0.233 |
| Llama 3.3 70B | 163 | 129 | 24 | 0.147 | 0.098–0.202 |
| Nemotron 120B | 163 | 163 | 45 | 0.276 | 0.215–0.344 |
| GPT-4.1 | 163 | 163 | 31 | 0.190 | 0.135–0.252 |
| GPT-4.1-mini | 163 | 163 | 31 | 0.190 | 0.135–0.252 |
| GPT-4o-mini | 163 | 154 | 29 | 0.178 | 0.123–0.239 |
| GPT-5.6 Luna | 163 | 0 | 0 | 0.000 | 0.000–0.000 |
| Qwen3-0.6B v3 | 163 | 163 | 33 | 0.202 | 0.141–0.270 |
| Qwen3-0.6B v5 | 163 | 163 | 32 | 0.196 | 0.141–0.258 |
| Qwen3-1.7B v3 | 163 | 163 | 31 | 0.190 | 0.135–0.252 |
| Qwen3-1.7B v5 | 163 | 163 | 40 | 0.245 | 0.184–0.319 |
| Qwen2.5 72B | 163 | 135 | 27 | 0.166 | 0.110–0.227 |
| Qwen3 235B | 163 | 158 | 30 | 0.184 | 0.123–0.245 |

#### Table S2c MB-2003

| Model | N | Parsed | Correct | Accuracy | 95% interval |
|---|---|---|---|---|---|
| Nova Pro | 300 | 200 | 99 | 0.330 | 0.241–0.423 |
| Claude Sonnet 4.5 | 300 | 299 | 183 | 0.610 | 0.503–0.708 |
| Mercury-2.5 | 300 | 268 | 131 | 0.437 | 0.330–0.541 |
| Llama 3.3 70B | 300 | 84 | 46 | 0.153 | 0.091–0.231 |
| GPT-4.1 | 300 | 152 | 83 | 0.277 | 0.192–0.369 |
| GPT-4.1-mini | 300 | 201 | 111 | 0.370 | 0.269–0.474 |
| GPT-4o-mini | 300 | 103 | 56 | 0.187 | 0.109–0.269 |
| GPT-5.6 Luna | 300 | 294 | 171 | 0.570 | 0.464–0.671 |
| Qwen3-0.6B v3 | 300 | 300 | 178 | 0.593 | 0.491–0.693 |
| Qwen3-0.6B v5 | 300 | 300 | 140 | 0.467 | 0.355–0.578 |
| Qwen3-1.7B v3 | 300 | 300 | 141 | 0.470 | 0.366–0.581 |
| Qwen3-1.7B v5 | 300 | 300 | 153 | 0.510 | 0.400–0.621 |
| Qwen2.5 72B | 300 | 146 | 76 | 0.253 | 0.173–0.343 |
| Qwen3 235B | 300 | 27 | 12 | 0.040 | 0.009–0.081 |
| DeepSeek Chat | 258 | 209 | 108 | 0.419 | 0.311–0.530 |
| Nemotron 120B | 258 | 258 | 129 | 0.500 | 0.388–0.615 |
| Gemini 2.5 Pro | 414 | 5 | 2 | 0.005 | 0.000–0.020 |

Pairwise rows belong to separate 300-, 258- or 414-item manifests and must not be ranked across those groups.

#### Table S2d MB-2005

| Model | N | Parsed | In domain | Conditional MAE | In-domain MAE | Failure-aware MAE |
|---|---|---|---|---|---|---|
| Nova Pro | 162 | 162 | 155 | 6.79e+03 | 1.710 | 1.809 |
| Claude Sonnet 4.5 | 162 | 162 | 136 | 2.77e+04 | 1.121 | 1.583 |
| DeepSeek Chat | 162 | 162 | 162 | 1.778 | 1.778 | 1.778 |
| Gemini 2.5 Pro | 162 | 0 | 0 | — | — | 4.000 |
| Mercury-2.5 | 162 | 109 | 102 | 3.08e+03 | 1.604 | 2.491 |
| Llama 3.3 70B | 162 | 150 | 150 | 1.517 | 1.517 | 1.701 |
| Nemotron 120B | 162 | 162 | 131 | 2.074 | 1.748 | 2.179 |
| GPT-4.1 | 162 | 162 | 162 | 1.802 | 1.802 | 1.802 |
| GPT-4.1-mini | 162 | 162 | 162 | 1.790 | 1.790 | 1.790 |
| GPT-4o-mini | 162 | 151 | 149 | 2.73e+03 | 1.611 | 1.802 |
| GPT-5.6 Luna | 162 | 20 | 20 | 1.800 | 1.800 | 3.728 |
| Qwen3-0.6B v3 | 162 | 162 | 162 | 1.481 | 1.481 | 1.481 |
| Qwen3-0.6B v5 | 162 | 162 | 140 | 2.06e+12 | 1.264 | 1.636 |
| Qwen3-1.7B v3 | 162 | 162 | 161 | 1.426 | 1.429 | 1.444 |
| Qwen3-1.7B v5 | 162 | 162 | 162 | 1.142 | 1.142 | 1.142 |
| Qwen2.5 72B | 162 | 162 | 162 | 1.617 | 1.617 | 1.617 |
| Qwen3 235B | 162 | 162 | 157 | 14.210 | 1.707 | 1.778 |

#### Table S2e MB-2006

| Model | N | Parsed | In domain | Conditional MAE | In-domain MAE | Failure-aware MAE |
|---|---|---|---|---|---|---|
| Nova Pro | 163 | 152 | 143 | 9.48e+03 | 3.734 | 4.258 |
| Claude Sonnet 4.5 | 163 | 163 | 163 | 3.828 | 3.828 | 3.828 |
| DeepSeek Chat | 163 | 157 | 157 | 3.994 | 3.994 | 4.141 |
| Gemini 2.5 Pro | 163 | 0 | 0 | — | — | 8.000 |
| Mercury-2.5 | 163 | 100 | 97 | 5.981 | 4.088 | 5.672 |
| Llama 3.3 70B | 163 | 154 | 152 | 2.46e+03 | 4.138 | 4.399 |
| Nemotron 120B | 163 | 162 | 155 | 3.833 | 3.690 | 3.902 |
| GPT-4.1 | 163 | 163 | 163 | 4.061 | 4.061 | 4.061 |
| GPT-4.1-mini | 163 | 163 | 163 | 4.215 | 4.215 | 4.215 |
| GPT-4o-mini | 163 | 162 | 162 | 4.111 | 4.111 | 4.135 |
| GPT-5.6 Luna | 163 | 0 | 0 | — | — | 8.000 |
| Qwen3-0.6B v3 | 163 | 163 | 155 | 6.13e+26 | 3.529 | 3.748 |
| Qwen3-0.6B v5 | 163 | 163 | 157 | 2.13e+02 | 2.968 | 3.153 |
| Qwen3-1.7B v3 | 163 | 163 | 162 | 2.221 | 2.154 | 2.190 |
| Qwen3-1.7B v5 | 163 | 163 | 158 | 6.13e+26 | 1.487 | 1.687 |
| Qwen2.5 72B | 163 | 163 | 163 | 4.067 | 4.067 | 4.067 |
| Qwen3 235B | 163 | 163 | 158 | 28.374 | 3.848 | 3.975 |

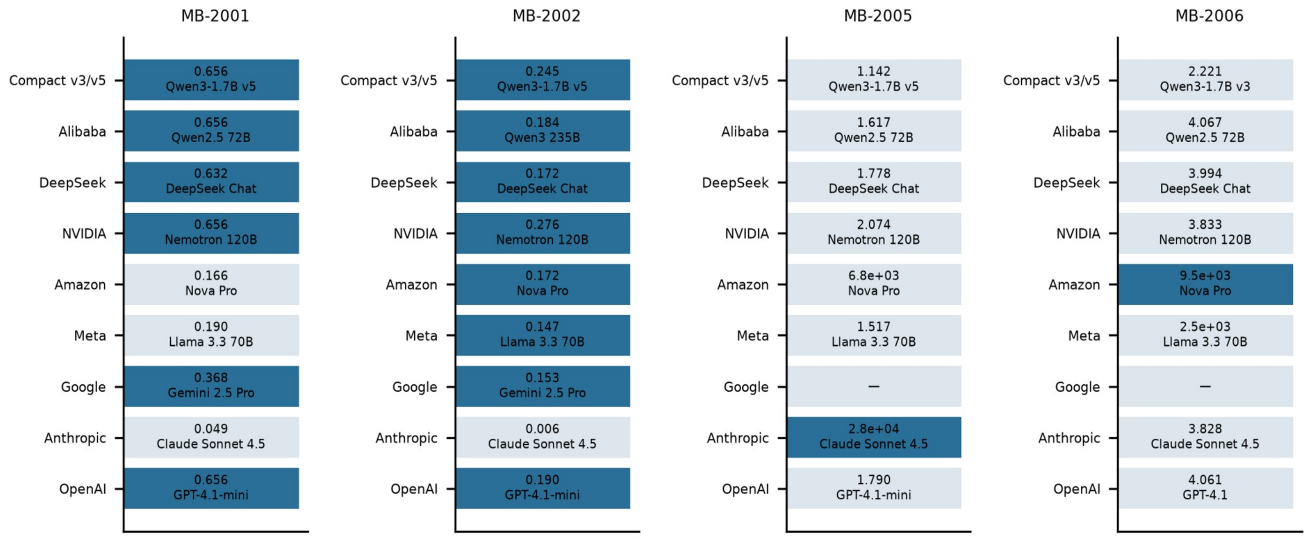


**Figure S1.** Best eligible model within each developer/group on the four aligned single-sample tissue tasks: binary advanced fibrosis (MB-2001), five-class fibrosis stage (MB-2002), numeric fibrosis stage (MB-2005), and numeric NAFLD Activity Score (MB-2006). Model selection is descriptive and performed on these evaluation sets. Pairwise results are omitted because manifests differ. Values are accuracy for classification and conditional MAE for regression; extreme values indicate numeric-format failures.

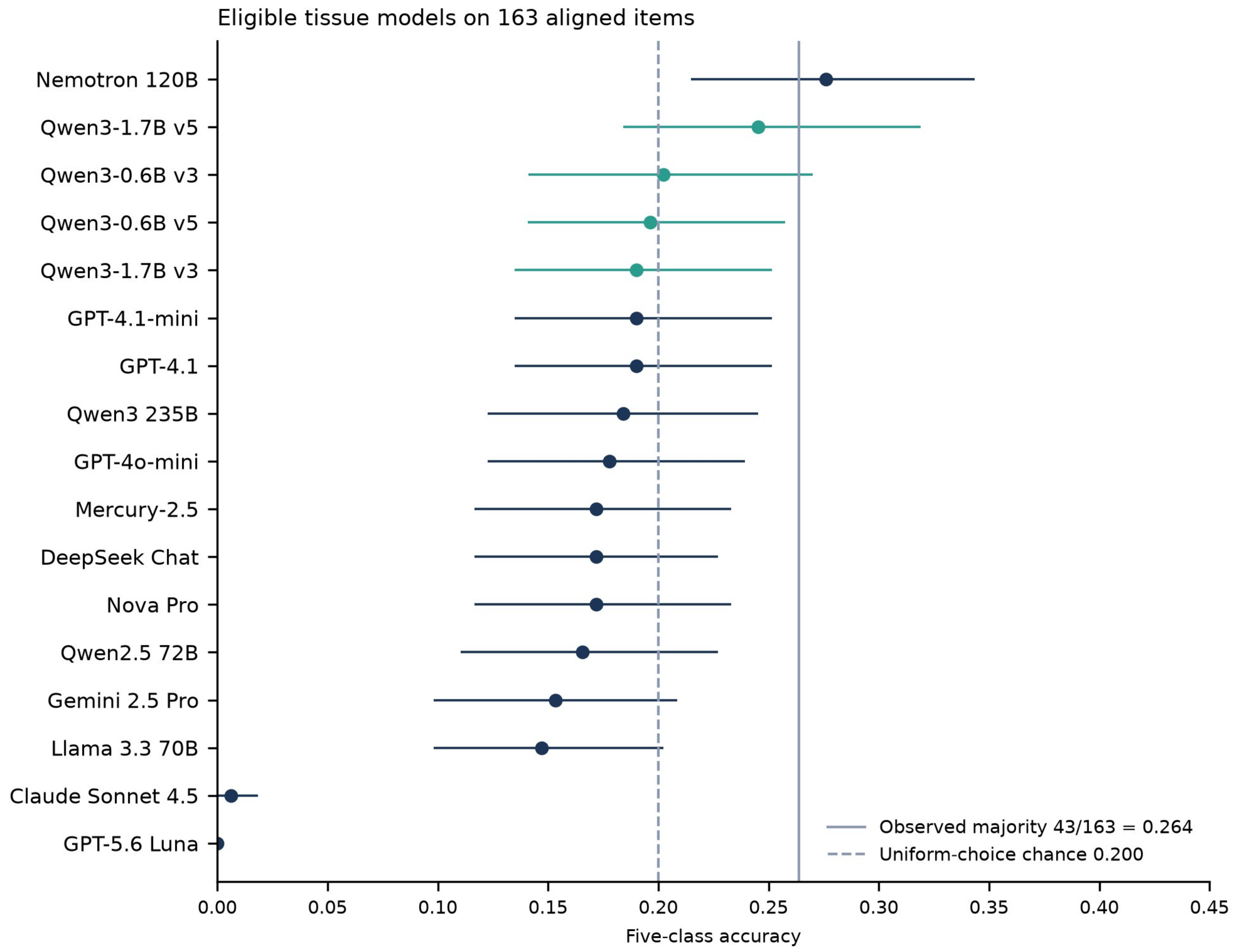


**Figure S2.** Five-class accuracy for 17 eligible tissue models on 163 aligned items. The solid line is observed majority 43/163 = 0.264 and the dashed line uniform-choice chance 0.200. Historical blood-proxy and plasma panels are excluded because they were not re-derived on the authoritative path.

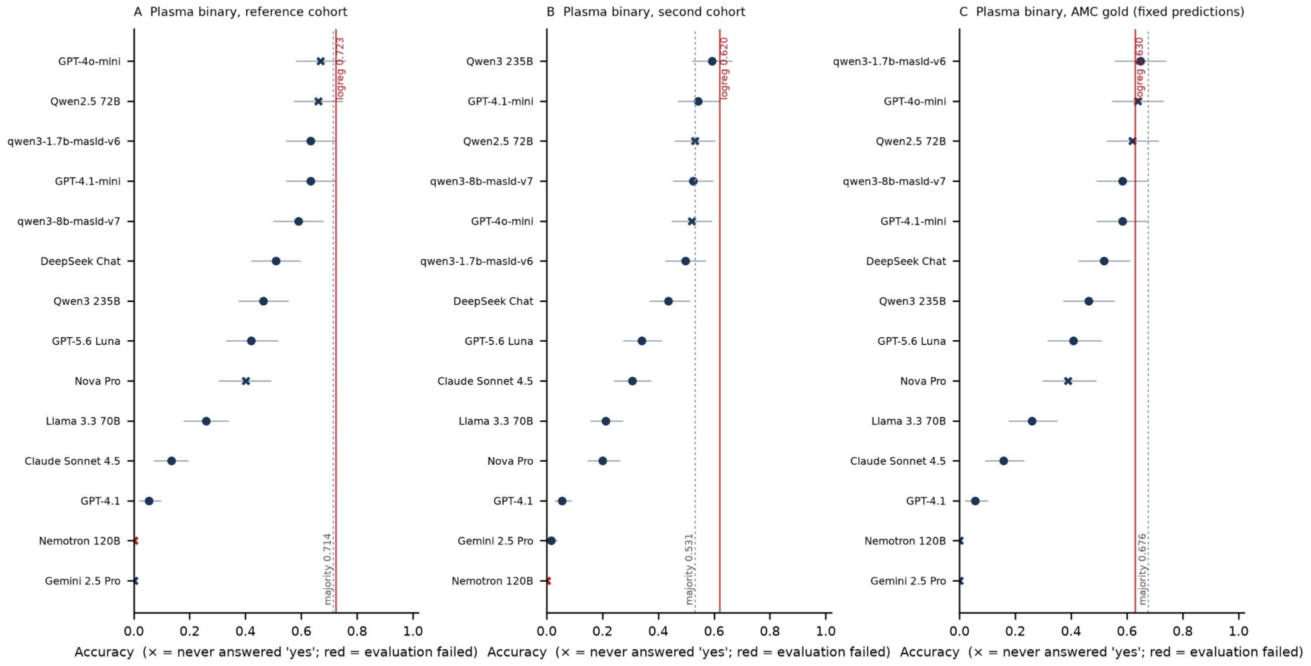


**Figure S3. Plasma binary results on reference-cohort, second-cohort and AMC-reader labels. Dashed majority lines are 0.714, 0.531 and 0.676. Red classical references are 0.723, 0.620 and 0.630. AMC uses fixed reference-cohort model predictions on 108 samples; it is a reader sensitivity, not an independent external cohort.**

#### Plasma cfRNA cohorts and preprocessing

##### Reference cohort

112 samples; biopsy-stage reference
Binary majority 80/112 = 0.714
Frozen top-200 expression panel

##### Reader subset

108 jointly staged reference-cohort samples
Exact agreement 56/108
Unweighted kappa 0.400

##### Second cfRNA cohort

179 patients; clinical-consensus reference
42-patient biopsy subpanel
Binary majority 95/179 = 0.531

##### Frozen representation

Gene-wise scaling across samples
Corpus-wide panel and moments
Transductive classical evaluation

##### Training-only sensitivities

Panel selected within each training fold
Training-only gene-wise moments OR
Per-sample standardisation

##### External five-gene transfer

Align genes by Ensembl identifier
Freeze reference-cohort-derived moments
Paired comparison remains inconclusive

The AMC reader subset is not an independent external cohort.

**Figure S4.** Cohorts and preprocessing contracts for cfRNA. The frozen representation uses gene-wise scaling across samples. Training-only sensitivities and external transfer are distinct. Reader agreement is based on 108 jointly staged samples.

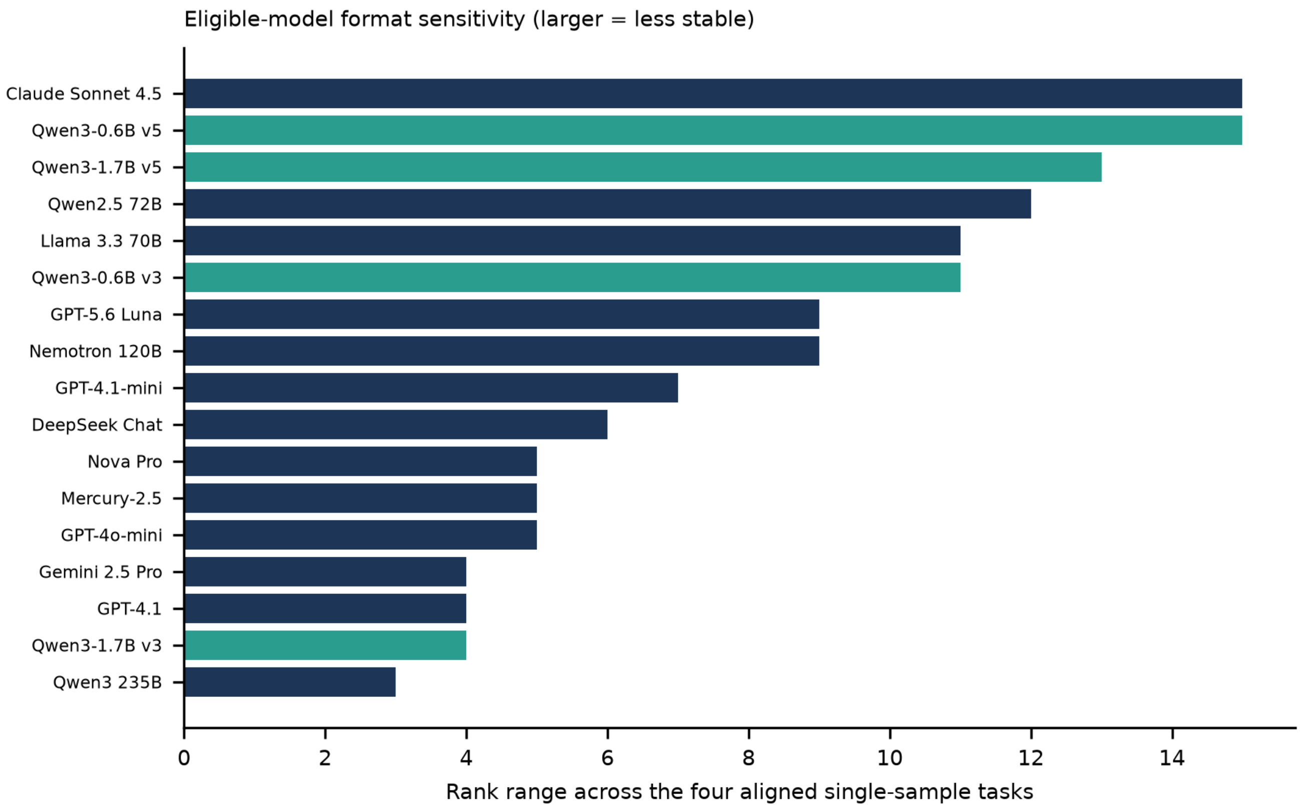


**Figure S5.** Within-model rank range over the four aligned single-sample tissue tasks. Pairwise ranks are excluded. Larger ranges indicate greater descriptive variation across task formats; they do not isolate format effects.

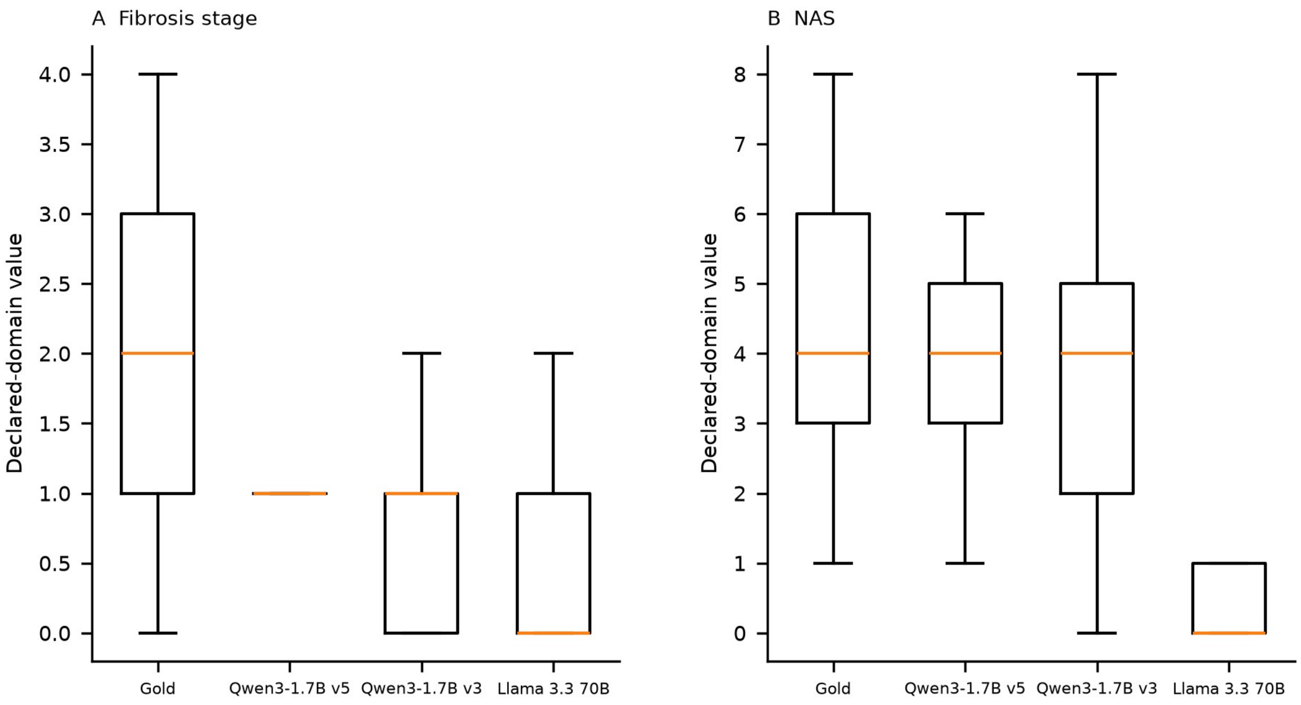


**Figure S6.** Gold and predicted distributions for selected eligible tissue models. Predictions are restricted to the declared domains (fibrosis 0–4; NAS 0–8). Boxes show interquartile ranges and median; whiskers follow the plotting library default and outliers are hidden. These parsing- and domain-selected distributions do not replace whole-set failure-aware results.

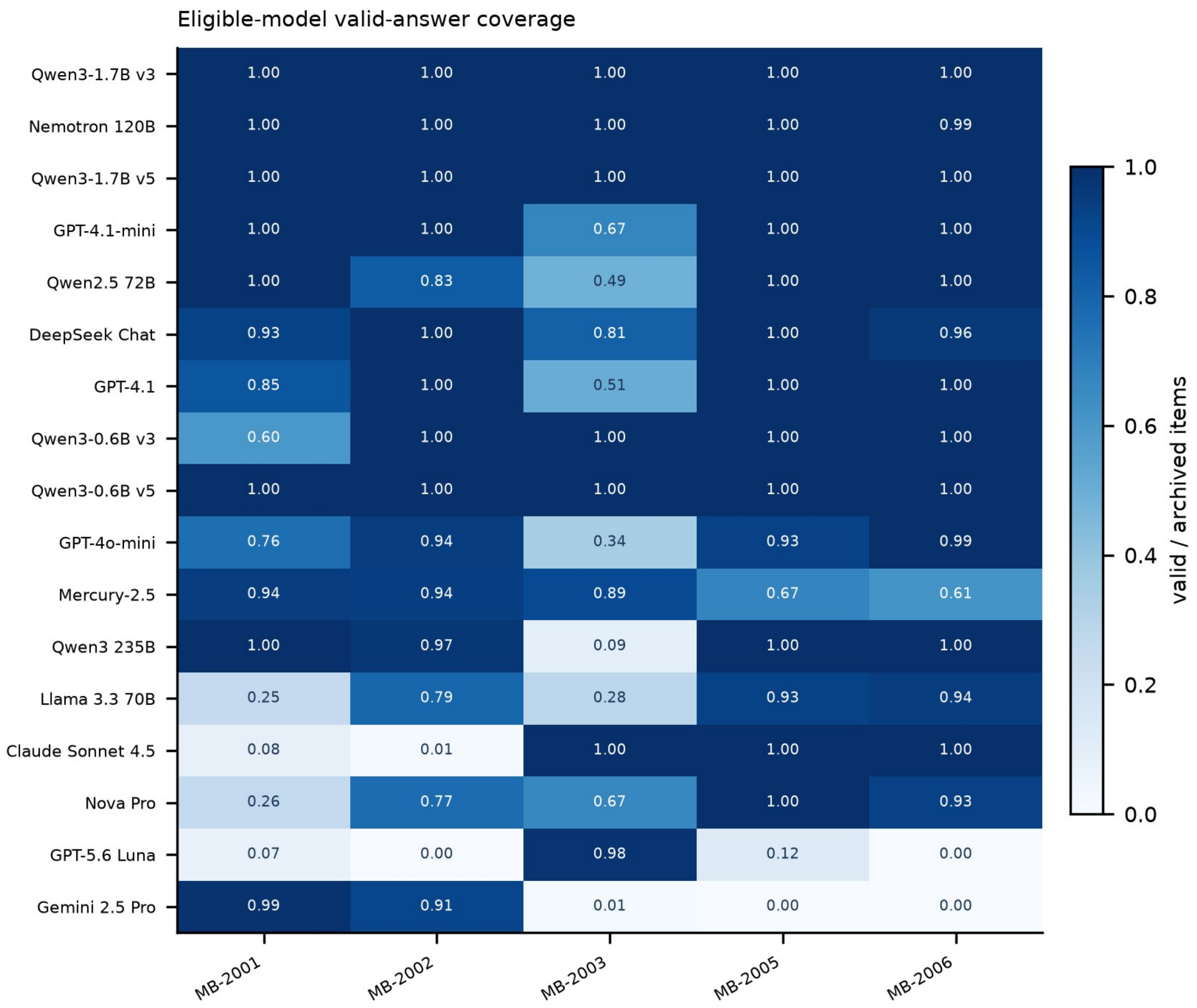


**Figure S7.** Parsed-answer coverage on aligned single-sample tasks and the respective pairwise manifests. Values are valid/archived items. Coverage is distinct from correctness and, for regression, from domain validity.

#### Distinct evaluation quantities

##### Resubstitution
Model evaluated on its training data
Reference-cohort five-gene accuracy 0.759

##### Conditional classifier cross-validation
Panel and scaling held fixed
Reference-cohort five-gene accuracy 0.714

##### Fully nested cross-validation
Panel, ranking and scaling re-derived
Top-5 reference-cohort accuracy 0.723

##### External transfer
Reference-cohort-fixed genes and transformation
External accuracy 0.615

##### Marginal-interval gate
Accuracy interval versus observed majority
Not a paired superiority comparison

##### Paired external comparison
51 model wins; 36 baseline wins
Exact McNemar $p = 0.133$

These quantities answer different questions; point estimates alone do not establish an advantage.

**Figure S8.** Interpretation of distinct cfRNA validation quantities. Resubstitution, fixed-panel classifier cross-validation, fully nested cross-validation and external transfer evaluate different quantities. The external paired comparison does not establish superiority.

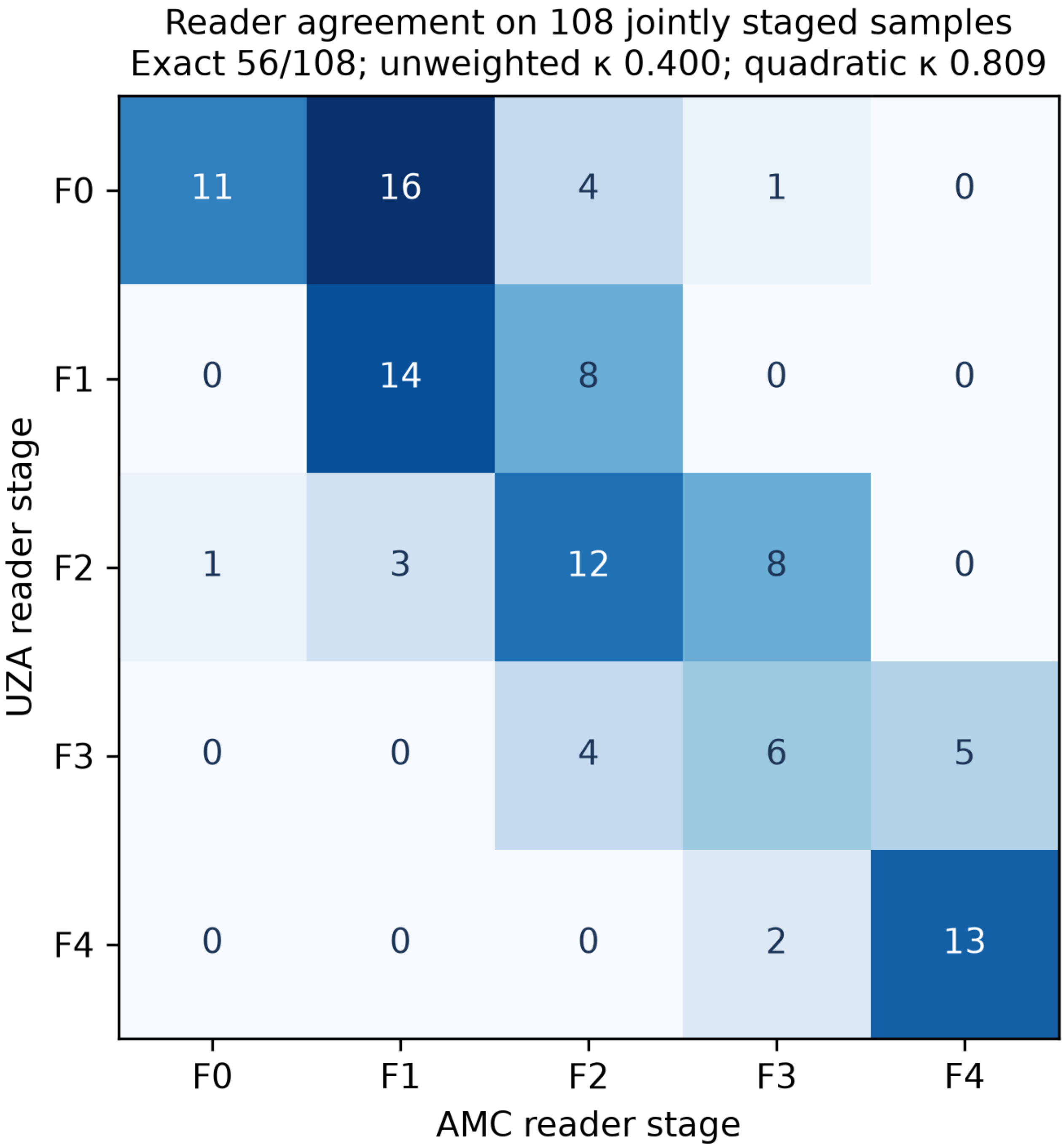


**Figure S9.** Confusion matrix between UZA and AMC biopsy readings for 108 jointly staged samples. Exact agreement is 56/108, unweighted κ = 0.400 and quadratic-weighted κ = 0.809. The aggregate matrix is reproduced from the corrected reader-agreement figure; no patient-level identifiers are included.
